# Probing the force-from-lipid mechanism with synthetic polymers

**DOI:** 10.1101/2022.05.20.492859

**Authors:** Miranda L. Jacobs, Jan Steinkühler, Audra Lemley, Megan J. Larmore, Taylor F. Gunnels, Leo C. T. Ng, Stephanie M. Cologna, Paul G. DeCaen, Neha P. Kamat

## Abstract

A central feature of mechanotransduction is the ability of mechanosensitive channels to respond to mechanical stimuli from the surrounding lipid bilayer. Accordingly, the mechanical properties of membranes should play an important role in modulating force transmission to embedded channels, yet the nature of this relationship remains unclear for a wide class of mechanosensitive channels across prokaryotic and eukaryotic systems. Here, we use a synthetic amphiphile to modulate the membrane mechanical properties of cell-derived vesicles and probe channel activation. Using precise membrane mechanical characterization approaches that have rarely been used in conjunction with electrophysiology techniques, we directly characterize three membrane properties and the activation threshold of the *E. coli* mechanosensitive channel of large conductance (MscL). Our study reveals that decreases in the membrane area expansion modulus, K_A_, and bending rigidity, k_c_, correlate with increases in the pressure required to activate MscL and that this effect is reproducible with the mammalian channel, TREK-1. MD simulations demonstrate that polymer-mediated changes in interfacial tension is the best mechanism to describe these experimental results. Together, our results bolster the *force-from-lipids* mechanism by demonstrating the generality of the relationship between changes in specific membrane mechanical properties and the gating pressure of MscL and TREK-1. In addition, our results reveal the mechanical mechanism by which membrane amphiphiles alter the activity and sensitivity of mechanosensitive channels through changes in long-range force transmission.

## Introduction

Mechanosensitive channels are a class of membrane proteins which are found in all domains of life and play important roles ranging from cellular volume regulation to touch and hearing sensation (Brohawn et al., 2019; Cahalan et al., 2015; Cox et al., 2019; Dave et al., 2021; del Mármol et al., 2018; Procko et al., 2021; Sun et al., 2019; Vásquez et al., 2014; Y. Wang et al., 2021). Many mechanosensitive channels are thought to transduce mechanical forces into biochemical signals by opening or closing in response to tension in the lipid bilayer, referred to as the *force-from-lipids* mechanism (Guo and MacKinnon, 2017; Martinac et al., 2020; Reddy et al., 2019; Ridone et al., 2018; Teng et al., 2015). Through this mechanism, mechanosensitive channels are proposed to be affected by long-range mechanical forces in the membrane that affect the bilayer lateral pressure profile (Anishkin et al., 2014; Bruno et al., 2007; Cordero-Morales and Vásquez, 2018; Pliotas and Naismith, 2017). As membrane tension can propagate distances of tens of microns within seconds in motile cells and axons (Houk et al., 2012; Shi et al., 2022), and the diffusion of tension relies directly on membrane properties (Cohen and Shi, 2020), lipid components that alter membrane mechanical properties should alter force transmission within a membrane and impact the activation of embedded mechanosensitive channels (Johnson, 2015). Toward this idea, others have demonstrated the relationship between membrane composition and mechanosensitive channel function and have shown that changes in membrane composition may modulate force propagation to embedded channels (Brohawn et al., 2014; Caires et al., 2017; Nakayama et al., 2018; Nomura et al., 2012; Ridone et al., 2018). Yet, because different amphiphiles have different effects on mechanosensitive channel activity, the mechanism of their action and the role of membrane mechanical properties on channel activity remains unclear.

To date, it has been difficult to directly resolve the role of membrane mechanical properties in mechanosensitive channel activation for two key reasons. The first is that certain membrane mechanical properties are difficult to experimentally measure. The second is that it can be difficult to significantly change specific membrane properties using lipids alone to isolate or probe their impact on channel activation. Toward the former, a variety of studies have reported that the alteration of membrane properties, through changes in membrane composition, affects mechanosensitive function (Anishkin et al., 2014; Balleza, 2012; Bruno et al., 2007; Caires et al., 2017; Lundbæk et al., 2010, 2004; Ridone et al., 2020, 2018; Romero et al., 2019; Xue et al., 2020). Through these studies, the impact of various membrane properties such as area expansion modulus (K_A_), the extent to which the membrane stretches in response to tension, bending stiffness (k_c_), the energy required to bend and smooth thermal fluctuations in a membrane, fluidity, a measure of viscosity or the lateral diffusion of lipids in a membrane, and hydrophobic thickness, the length of the hydrophobic core of the membrane, have been proposed to play a role in mechanosensitive channel behavior (Caires et al., 2017; Nakayama et al., 2018; Ridone et al., 2018; Romero et al., 2019). However, an important knowledge gap is that most of these studies have relied on published estimates of membrane properties or used methods which measure the bulk property of cell membranes, which includes contributions from cytoskeletal mechanics. To better understand how membrane properties affect mechanosensitive channel activation, it would be useful to experimentally probe channel activity in plasma membranes devoid of a cytoskeleton, for example, by using cell-derived, giant plasma membrane vesicles (GPMVs) as a model system. Toward the second reason, the direct role of certain membrane mechanical properties on mechanosensitive channel behavior has been difficult to discern in the presence of purely natural lipids. For example, phospholipids, which are the most abundant class of lipids in living systems, impart only small changes to the area expansion modulus of membranes (Rawicz et al., 2000)— a physical property which is expected to play a significant role in the activation of channels regulated by a *force-from-lipids* mechanism (Wiggins and Phillips, 2005). The effect of such small changes on mechanosensitive channel behavior is often difficult to detect or distinguish from changes in other membrane properties. However, non-natural amphiphiles, such as the plastics family referred to as poloxamers, can often alter membrane properties to a greater extent than natural lipids (Bermudez et al., 2002; Jacobs et al., 2019; Rawicz et al., 2000). By using GPMVs and incorporating membrane amphiphiles that change the membrane properties in unique and amplified ways, we hypothesized that we could better distinguish the impact of specific properties on protein function.

Here, we experimentally reveal a membrane property that exerts significant influence on the gating of a mechanosensitive channel: interfacial tension. By studying the relationship between membrane properties associated with interfacial tension (e.g., membrane elasticity, and bending rigidity) and the pressure required to activate mechanosensitive channels expressed in mammalian U2OS cells, we validate a hypothesized but not yet confirmed role of interfacial tension in mechanosensitive channel activation. We first establish this relationship for *E. coli* MscL by using U2OS-derived GPMVs, poloxamer analogues, voltage-clamp electrophysiology studies, and a series of mechanical characterization techniques, and we subsequently extend these findings to the mammalian mechanosensitive channel TREK-1. Further, we performed MD simulations to probe the impact of poloxamers on bilayer properties and found that polymer-mediated modulation of interfacial tension is a likely mechanism driving changes in channel activity. Our results highlight a critical role of membrane amphiphiles in indirectly regulating channel activity via changes in the bulk biophysical properties of membranes, with important implications in defining a mechanism of force-sensing in various living systems from bacterial to neuronal mechanotransduction. As poloxamers are a class of nanoplastic, we comment on the potential impact of nanoplastics on cellular health through interaction with cellular membranes.

## Results

Toward the goal of understanding the relationship between membrane amphiphiles and membrane force-transmission to embedded proteins, we prepared channel-containing GPMVs which allow for both assessment of mechanosensitive ion channel activity and membrane mechanical properties (Steinkühler et al., 2021, 2019). GPMVs not only incorporate membrane proteins of interest, but also maintain biological membrane composition and have an overall larger size (>5 µm) required for mechanical characterization via micropipette aspiration, while excluding organelles and cytoskeletal structures that might interfere with measurements of membrane mechanical properties, as in live cells (Figure 1A) (Keller et al., 2009; Levental et al., 2016; Steinkühler et al., 2019). We chose to first focus on the activity of the well-studied Mechanosensitive Channel of Large Conductance (MscL) in GPMVs. This channel opens when the membrane is stretched to permit molecules to flow across the membrane; channel activity can therefore be measured using electrophysiology (Chang et al., 1998; Wang et al., 2014). Specifically, we used a gain of function MscL missense mutation, G22S, as opposed to MscL wild-type (WT), due to the lower activation threshold of the former which increased the number of GPMVs with pressure-gated MscL currents (Figure 1-figure supplement 1A) (Heureaux et al., 2014). We created U2OS cell lines stably expressing an MscL-green fluorescent protein (GFP) fusion protein (Figure 1-figure supplement 2), and induced GPMV formation by adding a vesiculation agent (Figure 1-figure supplement 3). We elected to use N-ethlymaleimide (NEM) rather than the more commonly used paraformaldehyde/dithiothreitol (PFA/DTT) formulation as the latter results in crosslinking of membrane components (Sezgin et al., 2012) and may prevent mechanosensitive channel activation (Figure 1-figure supplement 4). This method also enables the incorporation of membrane additives to modulate membrane mechanical properties (Figure 1-figure supplement 5). The addition of this agent resulted in GPMVs containing MscL visualized by GFP fluorescence and that were an appropriate size for micropipette aspiration measurements (Figure 1A).

**Figure 1.**
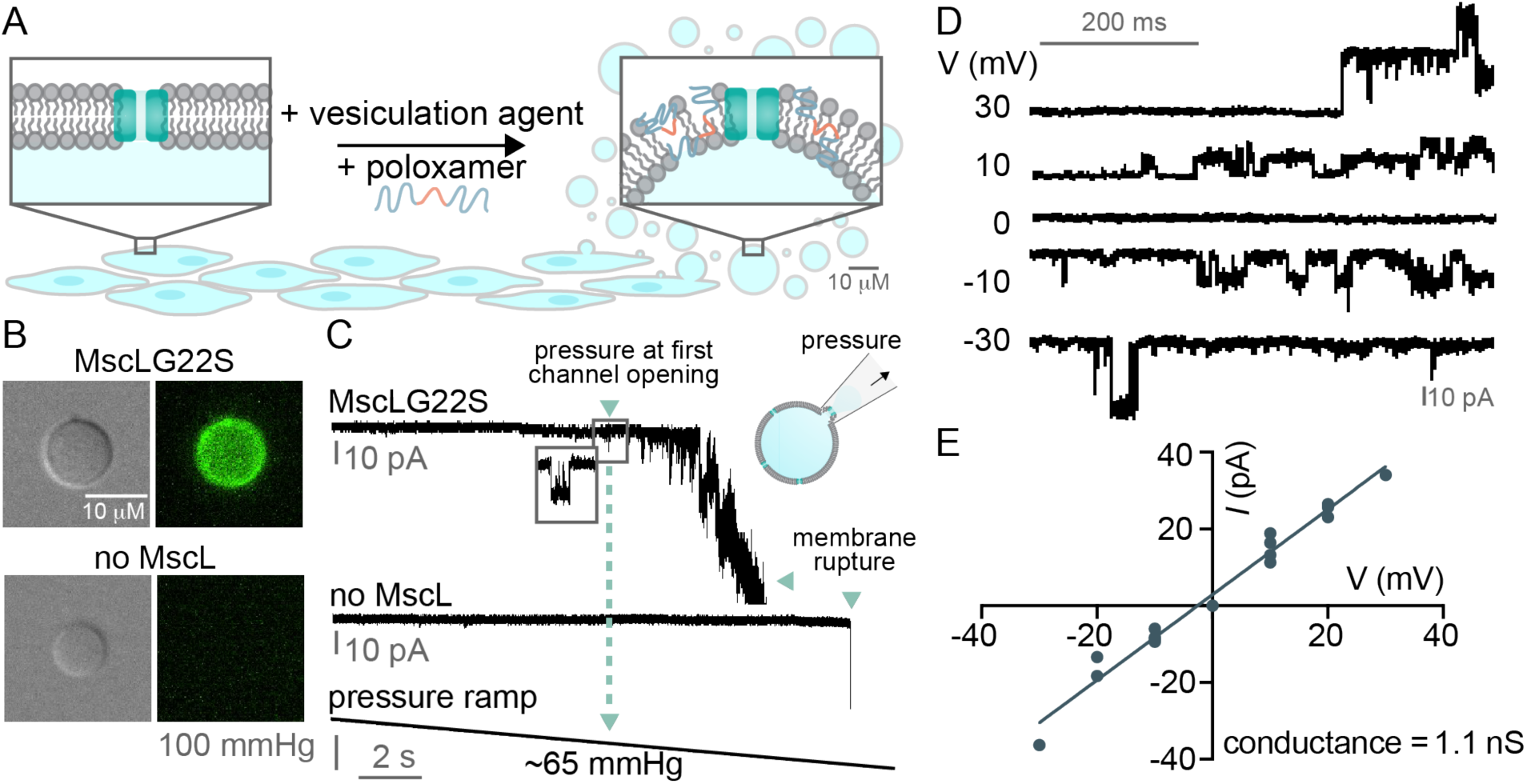
Patch-clamp electrophysiology of giant plasma membrane vesicles (GPMVs) measures MscL pressure sensitivity. A) Schematic representation of GPMV formation. Human bone osteosarcoma epithelial cells (U2OS) expressing MscLGFP were treated with N-ethylmaleimide (NEM) vesiculation agent to form GPMVs that contain MscLGFP. In certain cases, exogenous amphiphiles, such as poloxamers, were added to U2OS cells during the vesiculation process to generate GPMVs with modified membrane compositions. B) Microscopy images of GPMVs from non-transfected cells (bottom) or GPMVs from cells transfected with MscLG22S tagged with GFP (top). DIC images (left) and GFP epifluorescence (right) are shown. The scale bar is 10 μM. C) Representative electrophysiology traces of MscL pressure sensitivity over time. MscLG22S (top) recordings show MscL activation in response to pressure ramp (bottom). MscL activation sensitivity is defined as the pressure at which the first MscL channel activates (ex.∼65 mmHg for the shown GPMV). GPMVs from non-transfected cells (no MscL) exhibit no change in current in response to pressure (middle). Recordings were performed at -10 mV in 10 mM HEPES pH 7.3, 150 mM NaCl, 2 mM CaCl2. Pipette internal solution consists of 10 mM HEPES pH 7.3, 100 mM NaMES, 10 mM Na4BAPTA, 10 mM NaCl, 10 mM EGTA. D) Example MscLG22S single channel events recorded from GPMVs derived from U2OS cells at different voltages. E) Resulting single channel current-voltage relationship used in determination of the MscLG22S conductance. The current amplitude at each voltage (shown in D) is plotted and the slope of the linear fit was used to determine channel conductance. MscLG22S conductance is 1.113 nS (95% confidence 1.020 to 1.207 nS), n = 17 individual recordings at various holding potentials.

We next characterized our MscL-containing, GPMV system to confirm it retained key features of MscL function. We first validated the presence of MscLG22S in GPMVs via fluorescence microscopy using a C-terminal GFP tag, which was observable in our GPMV membranes (Figure 1B). Following this confirmation, we assessed MscLG22S activation sensitivity using patch-clamp electrophysiology techniques with an integrated pressure controller and quantified MscLG22S sensitivity using the pressure at first channel opening (Figure 1-figure supplement 6) (Nomura et al., 2012). We applied a pressure ramp that reached suction pressures above 160 mmHg and observed pressure-induced channel activation in our GPMVs that contained MscLG22S, while no currents were observed in our control GPMVs which lacked MscLG22S (Figure 1C, Figure 1-figure supplement 7). We measured channel conductance by quantifying the current through individual MscLG22S channels under different voltages (Figure 1D). We found the conductance of *E. coli* MscLG22S and MscLWT to be ∼1 nS in GPMVs (Figure 1E, Figure 1-figure supplement 1B), which is similar to published values of MscL conductance in mammalian cells [∼2 nS] (Doerner et al., 2012) and bacterial membranes [∼ 3 nS] (Sukharev et al., 1999, 1994). In addition, we confirmed that MscLG22S was sensitive to membrane composition in our GPMV platform through the addition of a small inverted cone shaped amphiphile, lysophosphatidylcholine (LPC) as described previously (Figure 1-figure supplement 8) (Nomura et al., 2012). These observations confirmed that the stochastic channel currents we observed were due to pressure-induced MscLG22S activation and that MscLG22S is sensitive to membrane composition in GPMVs, similar to other membrane platforms. This is also, to the best of our knowledge, the first demonstration that MscL activity can be recapitulated in GPMVs, further supporting a bilayer-mediated gating of the channel as originally shown in liposomes (Sukharev et al., 1994).

### Poloxamer P124 reduces the pressure-sensitivity of MscL

After characterizing the behavior of MscLG22S in unmodified GPMVs, we next set out to modify the membrane composition to determine the effect of various membrane mechanical properties on MscLG22S activation sensitivity. To modulate membrane properties, we used a poloxamer, which is an amphiphilic triblock copolymer that can integrate into bilayer membranes (Figure 2A) (Großkopf et al., 2021). We included increasing amounts of poloxamer 124 (P124) in the buffer during GPMV formation (Figure 1A) and measured MscLG22S activation pressure in the vesicles using patch-clamp electrophysiology. To confirm that poloxamers can integrate into the membrane, we used MALDI-MS and observed that two types of selected poloxamers, P124 and P188, integrate into membranes (Figure 2-figure supplement 1). Using microscopy, we found that poloxamer addition in certain cases increases GPMV diameter and surface area (Figure 2-figure supplement 2). In addition, we found that P124 increases GPMV stability as determined by resistance to rupture, consistent with previous studies (Figure 2-figure supplement 3) (Calori et al., 2022). Together, these results suggest that poloxamer readily integrates into GPMV membranes.

**Figure 2.**
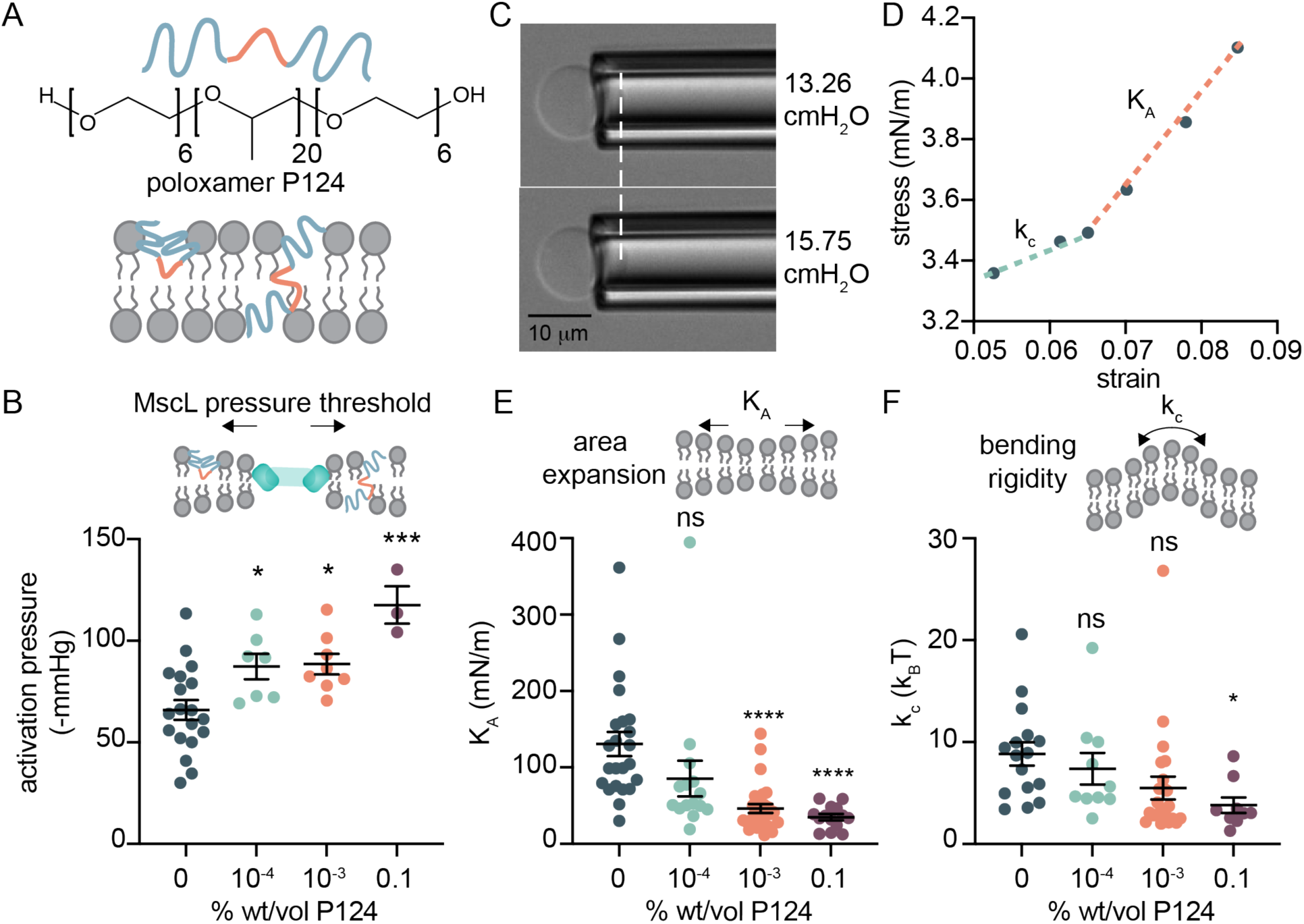
Poloxamer P124 decreases MscL sensitivity and modulates K_A_ and k_c_. A) Schematic and chemical structure of poloxamer P124. Poloxamer may be found in a hairpin or transverse conformation within the membrane. B) MscLG22S activation is desensitized in the presence of P124. MscLG22S activation is defined here as the pressure where the first MscLG22S channel is activated. The pressure at first channel opening was measured for GPMVs prepared in solutions containing 0 – 0.1% wt/vol P124, n ≥ 13 independent GPMVs were recorded. C) Images of micropipette aspiration of GPMVs measuring membrane mechanical properties. Micropipette aspiration is used to measure membrane areal changes in response to applied tension. The area expansion modulus (K_A_) and bending rigidity (k_c_) of GPMV membranes treated with poloxamer were measured. D) Sample aspiration curve from a micropipette aspiration experiment. A stress-strain curve was plotted to calculate K_A_ and k_c_ of GPMV membranes using micropipette aspiration techniques at high and low tension, respectively. E, F) K_A_ and k_c_ decrease in the presence of P124. GPMV K_A_ and k_c_ were measured using micropipette aspiration techniques after inclusion of P124 in the GPMV formation buffer. k_c_ and K_A_ were calculated at low- and high-pressure regimes, respectively, where K_A_ is corrected for relaxation of thermal undulations using mean k_c_. P124 % wt/vol is as listed, pH 7.3. Bath and pipette solution consist of 10 mM HEPES pH 7.3, 150 mM NaCl, 2 mM CaCl2 in micropipette aspiration studies. Mean activation pressure, K_A_, k_c_ are plotted with error bars that represent standard error of the mean. p-values in B, E, and F were generated by ANOVA using Dunnett’s multiple comparisons test compared to no poloxamer. **** p ≤ 0.0001, *** p ≤ 0.001, * p ≤ 0.05, non-significant (ns) p >0.05, n>10.

We found that MscLG22S required higher pressures to activate when GPMVs contained P124 and this response was monotonic with increasing poloxamer concentration, indicating that P124 decreased the sensitivity of MscLG22S (Figure 2B). To further evaluate MscLG22S sensitivity in the presence of P124, we quantified the percent of GPMVs that contained MscLG22S currents out of all recorded GPMVs which exhibited GFP fluorescence. In this quantification, we only monitored currents up to a maximum pressure above -80 mmHg, under which we would normally expect MscLG22S activation in a poloxamer-free membrane. We found in GPMVs with the highest buffer concentration of P124 (0.1% wt/vol), only ∼20% of GPMV recordings contained active MscLG22S currents, indicating that MscLG22S was not activating in ∼80% of GPMVs prepared in high concentrations of P124 (Figure 2-figure supplement 4). Together, these results suggest that the pressure sensitivity of MscLG22S is reduced in the presence of P124 and increases in P124 content eventually leads to a large fraction of channels failing to productively open.

We then investigated how P124 altered GPMV mechanical properties. Using micropipette aspiration techniques, we measured the area expansion modulus (K_A_) (Jacobs et al., 2021) and bending rigidity (k_c_) (Rawicz et al., 2000) of GPMVs treated with P124 for greater than 4 hours (Figure 2C, D; Figure 2-figure supplement 5). We found that membrane K_A_ and k_c_ decreased with increasing P124 concentration in the buffer (Figure 2E, F). We then measured P124 fluidity using fluorescence anisotropy techniques and did not observe a significant change in membrane fluidity in the presence of P124 (Figure 2-figure supplement 6).

We wondered how these changes in mechanical properties induced by P124 may affect mechanosensitive channel behavior. To assess this relationship, we plotted MscLG22S activation pressure as a function of each of these membrane mechanical property measurements and found that decreases in membrane K_A_ and k_c_ correlate with commensurate increases in MscLG22S activation pressure, r = -0.89 and -0.99, respectively (Figure 2-figure supplement 7). These results suggest that the reduced sensitivity of MscLG22S to pressure in the presence of P124 may be related to the alteration of specific membrane mechanical properties.

### Various poloxamers and a detergent reveal a correlation between K_A_, k_c_ and MscL activation sensitivity

To further explore the generality of the relationship between membrane mechanical properties and MscLG22S activation sensitivity, we used a chemically distinct detergent and an expanded set of poloxamers. We selected the detergent C12E8, which is known to decrease membrane K_A_ (Jacobs et al., 2019; Otten et al., 2000), and measured the effect of this molecule on MscLG22S activation sensitivity. Similar to our results in the presence of P124, C12E8 increased the pressure required to activate MscLG22S (Figure 3-figure supplement 1A) although C12E8 did not depress the percent of active channels to the same extent as P124 (Figure 3-figure supplement 1B). We then measured the mechanical properties of GPMVs treated with C12E8 and confirmed that this molecule has a softening effect on membranes, reducing the K_A_ and k_c_ (Figure 3-figure supplement 1C, D, E). We then measured the effect of other poloxamers with similar chemical structures but differing molecular weights on MscLG22S pressure sensitivity and membrane mechanical properties (Figure 1-figure supplement 5, Figure 3-figure supplement 2, 3). We plotted the activation pressure of MscLG22S as a function of K_A_, k_c_, and fluidity for each of the amphiphiles in which we measured both MscLG22S activation and membrane properties and found similar trends as when we evaluated these relationships for each amphiphile individually (Figure 3, Figure 3-figure supplement 4, Figure 2-figure supplement 7). We found that membrane K_A_ (r = -0.71) (Figure 3A) and k_c_ (r = -0.81) (Figure 3B) correlated with MscLG22S activation pressure, however, as we observed in the GPMVs containing P124, fluidity did not correlate well with MscLG22S activation pressure (r = 0.15) (Figure 3C). Taken together, these results confirm that MscLG22S behavior is sensitive to changes in membrane composition and properties where conditions with reduced K_A_ and k_c_ generally correlate with a reduction in the pressure sensitivity of MscLG22S (Figure 3D).

**Figure 3.**
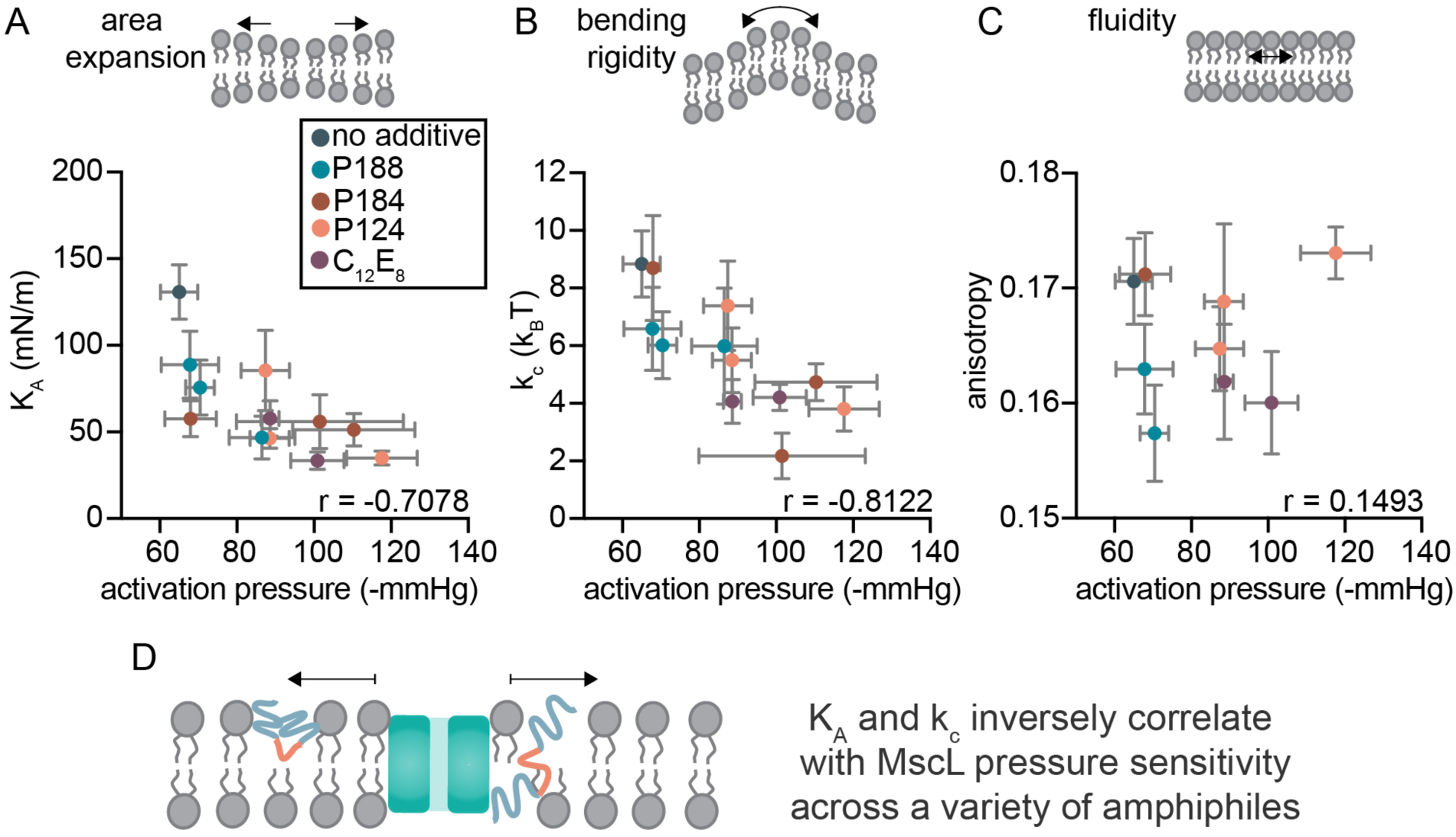
Membrane area expansion modulus and bending rigidity are good predictors for MscL pressure sensitivity. A, B, C) Membrane K_A_ and k_c_, but not fluidity, predict MscLG22S activation pressure. GPMVs were treated with various poloxamers or a detergent (C12E8) during formation and membrane properties (KA, k_c_, and fluidity) and MscLG22S pressure sensitivity were measured. Mean K_A_ (A), k_c_ (B), and fluidity (via anisotropy) (C) are reported as a function of MscLG22S activation pressure when treated with the same amphiphile at a given concentration. Decreased anisotropy indicates a relative increase in membrane fluidity. For all plots, Pearson’s r was used to determine correlation strength, n > 10. D) Summary of key correlations (KA and k_c_) between GPMV membrane properties and MscLG22S activation sensitivity across a variety of amphiphile additives.

### P124 does not alter MscL activity through pore occlusion or changes in membrane thickness but may function through reduction in interfacial tension

We sought to understand the mechanism behind amphiphile-induced changes in MscLG22S activation sensitivity, and we elected to rule out the possibility that P124 modulates MscLG22S through a mechanism other than altering membrane mechanical properties. We first hypothesized that P124 could modulate MscLG22S through pore occlusion (Figure 4A). As MscLG22S conductance depends directly on pore size when measured in identical buffer conditions, a decrease in conductance would be expected if P124 resides within the MscLG22S pore (Banerjee et al., 2013; Rokitskaya et al., 2017). We measured the conductance of MscLG22S in the presence of increasing amounts of P124 and found that MscLG22S conductance was not affected by P124 (Figure 4B, C). We included C12E8 as a control which is chemically distinct from poloxamer and would therefore not be expected to interact with MscLG22S in the same way and found that C12E8 also did not affect MscLG22S conductance. Together, these observations suggest that P124 and C12E8 do not reside within the MscLG22S pore while in the active conformation.

**Figure 4.**
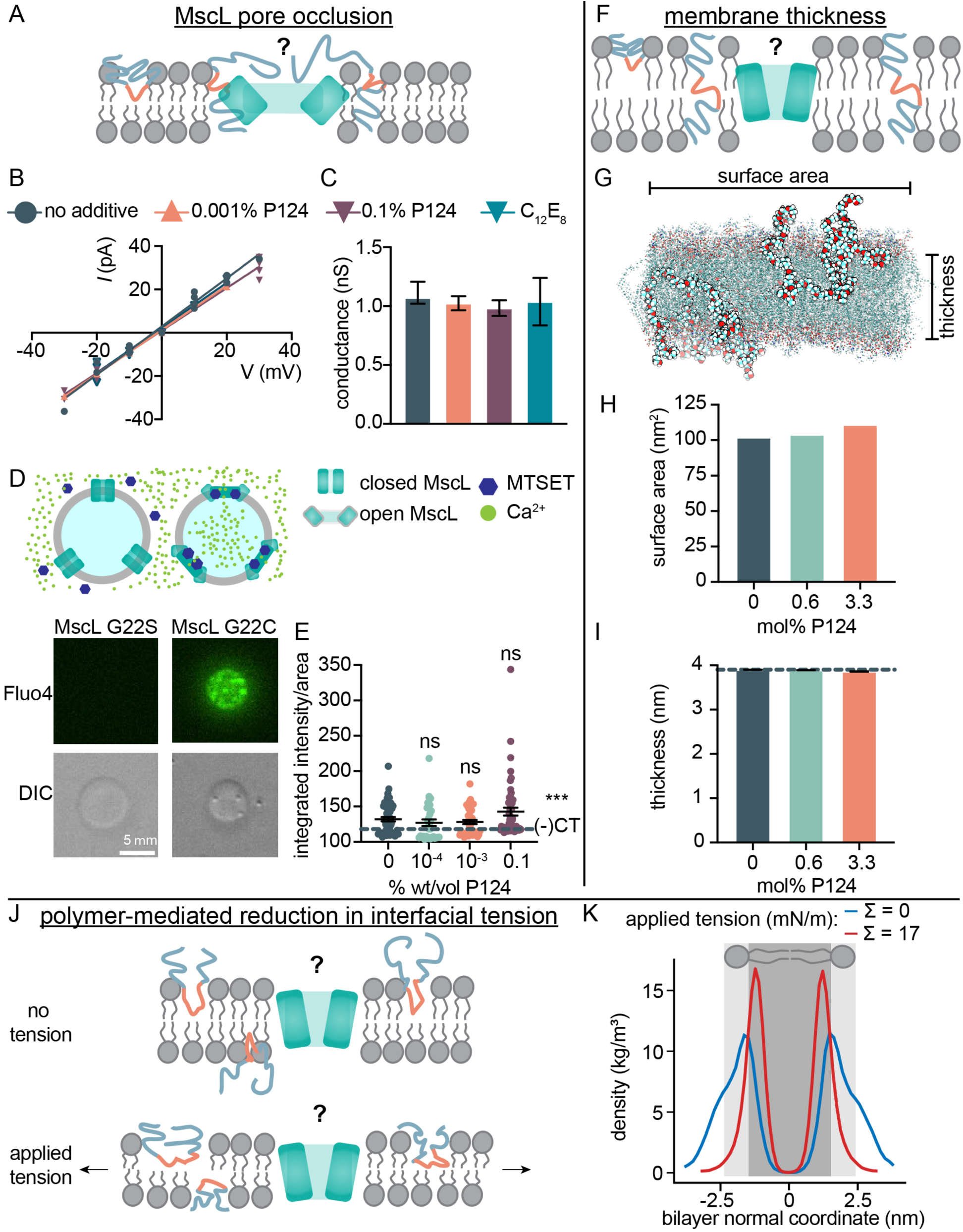
Poloxamer does not impact MscL through pore occlusion or membrane thickness changes but likely through a reduction in interfacial tension. A) Poloxamers could prevent MscLG22S activation through pore occlusion. B) MscLG22S pore size, as inferred by conductance calculations, does not change in the presence of P124. MscLG22S conductance in the presence of P124 is plotted; the slope of the linear fit was used to determine channel conductance. C12E8 serves as a non-polymeric control. C) MscLG22S conductance quantification in the presence of P124 and C12E8. Mean conductance and 95% confidence interval are plotted from the linear fit shown in (B). D) Chemical activation of MscLG22C. Fluo4 Ca^2+^ imaging of the internal space of the vesicle reveals that Ca^2+^ influx can be detected when MscLG22C is activated with MTSET but not in the presence of MscLG22S. DIC images show GPMV outline. E) P124 does not prevent MscL activation. GPMVs containing MscLG22C were formed in the presence of various P124 concentrations as indicated. Fluo4 fluorescence was quantified after GPMV incubation with MTSET and 5 mM Ca^2+^ and is plotted as integrated intensity / vesicle area. Mean is plotted with standard error off the mean, and the dashed horizontal line is the signal for MscLG22S in 0% P124 (negative control, (-)CT). *** p ≤ 0.001, non-significant (ns) p > 0.5, n > 30 vesicles per condition. p-values were generated by ANOVA using Dunnett’s multiple comparisons test compared to no poloxamer. F) Poloxamers may prevent MscLG22S activation by increasing membrane thickness. G) Schematic of coarse-grain molecular dynamics simulations of DOPC bilayers (translucent) containing P124 (opaque). H) Membrane surface area increases with increasing poloxamer concentration. I) P124 does not increase membrane thickness in molecular simulations. Error bars represent standard error of the mean, n = 3. J) Poloxamers may affect MscL channel opening by reducing interfacial tension in the membrane when under tension. K) Distribution of the PEO blocks normal to the lipid bilayer in P124-containing membranes as determined by coarse-grained molecular dynamics simulations. The normal coordinate at 0 is defined as the middle of the lipid bilayer, and the approximate location of the hydrophobic region is shaded dark gray and the hydrophilic region is shaded light gray. Under tension, the PEO groups adsorb to the bilayer, suggesting adsorption is a thermodynamically favorable process that lowers the interfacial tension.

To further address the possibility that P124 alters MscL activation through pore occlusion, we measured the propensity of P124 to alter activation of a chemically-inducible mutant of MscL that is insensitive to pressure. A mutation within the pore of MscL serves as a hydrophobic gate (G22C) which can be activated by the addition of [2-(trimethylammonium)ethyl] methanethiosulfonate bromide (MTSET), a chemical which forcibly hydrates the pore by introducing charge (Yoshimura et al., 2001). We hypothesized that if P124 occluded the MscL pore, MTSET would be unable to open the MscLG22C channel. To assess MscLG22C opening in GPMVs upon MTSET binding, we monitored Ca^2+^ influx via a fluorescent Ca^2+^ indicator, Fluo4. We first validated this assay by ensuring that internal vesicle fluorescence, indicating Ca^2+^ influx, increased only in the presence of the chemically activatable mutant of MscL G22C, and not the pressure sensitive mutant G22S, when MTSET was present (Figure 4D, Figure 4-figure supplement 1). We also confirmed MscLG22C’s lack of pressure sensitivity (Figure 4-figure supplement 2) (Levin and Blount, 2004). We then determined if MscLG22C chemical activation was altered by P124 addition by quantifying the internal fluorescence of GPMVs containing P124 in the presence of MTSET. This assay showed that P124 did not inhibit MscLG22C activation relative to the no P124 control (Figure 4E) and all conditions resulted in small but detectable increases in Ca^2+^-induced fluorescence relative to the MTSET-insensitive control. Taken together, these results suggest that P124 does not occlude the MscL pore.

Another potential mechanism by which MscLG22S activation could be altered in the presence of P124 is through increased membrane thickness (Figure 4F). MscL is known to be sensitive to membrane thickness where thinner membranes reduce the gating threshold of MscL (Ridone et al., 2018) and thicker membranes desensitize MscL (Nomura et al., 2012; Ridone et al., 2018). We do not expect membrane thickness to be increased in the presence of poloxamer based on our bending rigidity measurements, because the reduced k_c_ values suggest a decreased membrane thickness (Figure 3B). To further investigate this hypothesis, we performed molecular dynamics simulations to measure changes in membrane thickness, surface area, and K_A_ after P124 incubation (Figure 4G). Our simulations demonstrated that P124 inserts into the bilayer and this integration increases surface area (Figure 4H) and reduces membrane K_A_ (Figure 4-figure supplement 3) but does not alter membrane thickness significantly (Figure 4I).

To better understand how polymers affect the mechanical properties of our membranes, we simulated the effect of mechanical tension on P124-containing lipid membranes (Figure 4J, 4K).

At zero tension, the polyoxyethylene (PEO) hydrophilic groups of the polymer protrude deeply into the water phase, resulting in an extended, “PEGylated” lipid bilayer. The conformation of the polymer changes with applied tension. Specifically, we observed that PEO obtains a more compact conformation “focused” at the lipid-water interface under tension. The behavior reflects the fact that PEO has partially amphiphilic character (Israelachvili 1997), and the lipid bilayer prefers to adsorb PEO to minimize exposure to water molecules that might otherwise penetrate the hydrophobic region of the membrane exposed under tension. Spontaneous adsorption of PEO groups (i.e., a thermodynamically-favored process) implies that the water-bilayer interfacial tension is reduced by P124 when under increased mechanical tension. Since interfacial tension is a known driver of MscL activity (Melo et al., 2017; Ollila et al., 2011), these data suggest a potential mechanism for how P124 increases MscL’s opening pressure despite decreasing membrane elastic properties K_A_ and k_c_.

### TREK-1 activation sensitivity also correlates with K_A_ and k_c_ and supports the *force-from-lipids* gating of mechanosensitive channels

Finally, we sought to explore the generality of our findings with another mechanosensitive channel. We chose to measure the effect of P124 on the mouse potassium channel, TWIK-related K+ channel 1 (TREK-1), a mechanosensitive channel found in neurons that can be activated by *force-from-lipids* and is sensitive to changes in membrane composition (Brohawn et al., 2014). Using a similar expression scheme as for MscL, we created stable cell lines of TREK-1 tagged with mEGFP (Figure 5-figure supplement 1), which was retained in the membrane after GPMV formation (Figure 5-figure supplement 2). We observed TREK-1 single-channel currents in response to a pressure gradient using an integrated pressure controller. Similar currents were not observed in non-transfected cells, lacking the channel, and exposed to the same experimental conditions (Figure 5A, Figure 5-figure supplement 3). We measured TREK-1 conductance using voltage-clamp electrophysiology measuring the respective current through a single channel and observed an outward conductance of 140 pS, similar to the conductance of TREK-1 in cellular systems (Figure 5B) (Heurteaux et al., 2004; Xian Tao Li et al., 2006). The inward conductance was undetectable below +50 mV, suggesting a primarily outward rectifying channel, consistent with previous observations (Xian Tao Li et al., 2006). We next determined if TREK-1 activation pressure was altered by P124. We treated TREK-1-containing GPMVs with P124 and observed an increase in the pressure required to activate the first TREK-1 channel (Figure 5C). Again, the chemically distinct C12E8 amphiphile also increased the channel activation pressure (Figure 5C), similar to our results with MscL (Figure 2B, Figure 3-figure supplement 1A). Finally, we wondered if TREK-1 activation could be predicted by membrane properties. We plotted K_A_, k_c_, and fluidity as a function of TREK-1 activation pressure. Similar to MscL, we observed a strong correlation between TREK-1 activation pressure and K_A_ (r = -0.99) and k_c_ (r = -0.97) (Figure 5D, E). As we observed for MscL, there was not a strong correlation between membrane fluidity and TREK-1 activation pressure (r = -0.50) (Figure 5F). These results suggest that, as for MscL, a correlative relationship exists between membrane elastic properties, K_A_, and k_c_, and TREK-1 activation sensitivity. Taken together, our observations suggest that inclusion of non-natural amphiphiles can drive changes in membrane mechanical properties that are sufficient to alter mechanosensitive channel activation.

**Figure 5.**
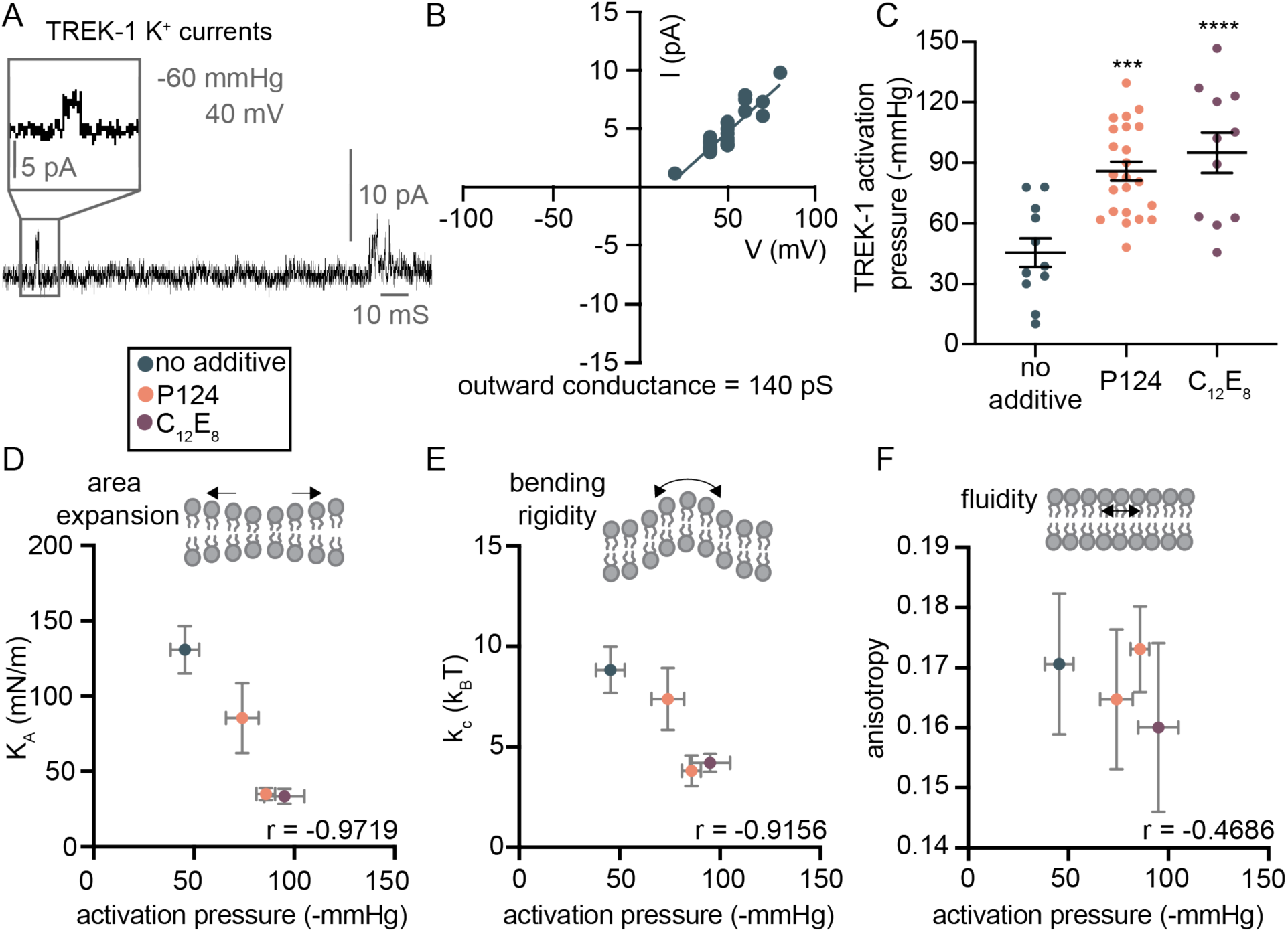
TREK-1 activation also correlates with membrane properties. A) Mouse TREK-1 single-channel potassium currents are observed in GPMVs in response to pressure. TREK-1 currents were measured using patch-clamp electrophysiology with an integrated pressure controller in asymmetric potassium conditions, pH 7.3. B) TREK-1 is outward rectifying with a conductance of 140 pS. TREK-1 current was quantified under various holding potentials. TREK-1 currents were larger under negative voltages and the slope of the I/V relationship under negative voltages was used to measure outward rectifying conductance. Individual electrophysiology measurements are plotted, n > 15. C) TREK-1 activation pressure is increased in the presence of a poloxamer (0.1% P124) and a detergent (1 μm C12E8). Mean activation pressure is plotted and dots represent individual measurements, error bars represent standard error of the mean. p-values were generated by ANOVA using Dunnett’s multiple comparisons test compared to no poloxamer. **** p ≤ 0.0001, *** p ≤ 0.001, n>10. D, E) Membrane K_A_ and k_c_ predict TREK-1 activation pressure. GPMVs were treated with P124 at two concentrations and C12E8 at the highest concentration and TREK-1 activation or membrane mechanical properties were measured. Mean K_A_ or k_c_ and TREK-1 activation pressure were plotted. F) Membrane fluidity does not predict TREK-1 pressure sensitivity. Mean membrane anisotropy and TREK-1 activation pressure were plotted. Decreased anisotropy indicates a relative increase in membrane fluidity. Pearson’s r in (D-F) was determined to measure correlation strength, error bars represent standard error of the mean, n > 10 for each measurement.

## Discussion

Through a series of micropipette aspiration and patch-clamp electrophysiology studies, we have determined the effect of various poloxamers and a chemically distinct detergent on membrane properties and mechanosensitive channel behavior. By measuring these membrane properties directly, we found that membrane K_A_ and k_c_ inversely correlate with activation sensitivity of a mechanosensitive MscL variant, while membrane fluidity does not to the same extent. We found that the amphiphiles we used did not appear to significantly change membrane thickness or occlude MscL pores, factors that would be expected to impact channel activity. Instead, polymer-mediated changes in MscL activation threshold are likely due to differences in bilayer mechanical properties. Finally, we determined that K_A_ and k_c_ also correlate to the behavior of more complex mechanosensitive channels in higher organisms by assessing the behavior of TREK-1 in response to changes in membrane properties. Our findings, together with molecular dynamics simulations, point to a specific mechanical property that influences pressure sensitivity of the studied channels: interfacial tension. By decreasing interfacial tension in the membrane, a greater force is required to activate MscL and TREK-1. To our knowledge, our study is the first to directly measure multiple membrane properties using GPMVs in conjunction with electrophysiology assessments to directly implicate interfacial tension, K_A_, and k_c_ in MscL and TREK-1 activation sensitivity.

Mechanosensitive channel activation sensitivity is an important feature of all mechanosensitive channels. Overactive mechanosensitive channels may lead to deleterious effects ranging from a loss in cell volume in bacteria or red blood cells (Cahalan et al., 2015; Levin and Blount, 2004) to persistent itch disease in higher organisms (LaMotte, 2016). Membrane composition has been increasingly shown to modulate the *force-from-lipids* activation of mechanosensitive channels and this effect is hypothesized to be due to changes in global membrane properties (Bruno et al., 2007). We recently showed that membrane amphiphiles can impact membrane properties in unpredictable ways (Jacobs et al., 2021) underlying the importance of directly measuring membrane properties when drawing conclusions about their role in a physiological outcome. While a number of studies discuss the effects of membrane properties on mechanosensitive channel activation (Nomura et al., 2012; Ridone et al., 2018; Xue et al., 2020), and there is increasing appreciation of the need to measure channel activity in response to applied membrane tension (Lewis and Grandl, 2015; Lüchtefeld et al., 2024), few have directly measured membrane properties (Caires et al., 2017; Nakayama et al., 2018; Romero et al., 2019) to determine their impact on channel function. Moreover, to the best of our knowledge no studies to date have used the classical micropipette aspiration technique (Kwok and Evans, 1981) in conjunction with patch-clamp electrophysiology to directly determine the relationship between membrane properties and mechanosensitive channel function. Accordingly, the body of research to date has left an unexplored gap between membrane mechanical properties and mechanosensitive channel behavior that we have sought to address. In the present study, we demonstrated that specific membrane properties modulate the *force-from-lipids* activation of two mechanosensitive channels.

How do our findings compare with the existing models of mechanosensitive channel activity? We start with a classic thermodynamical model of MscL opening, where the free energy difference, Δ*G*, between the open and closed states of MscL can be divided into three contributions (Ollila et al. 2011; Phillips et al. 2009; Wiggins and Phillips 2005). Here, Δ*G* refers to the free energy of opening the channel:

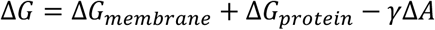

The term Δ*G*_*membrane*_refers to the energetic costs incurred from the elastic deformations of the lipid bilayer that are required to accommodate MscL, Δ*G*_*protein*_refers to the energy difference of the open and closed state of MscL (to include any changes in lipid-protein interactions), and −*γ*Δ*A* describes the impetus for channel activation which is the product of the protein surface area gained (Δ*A*) by opening the channel with applied tension (*γ*). We define ΔΔ*G*_*exp*_ as the free energy difference of channel opening between a lipid-only membrane and a modified membrane. We reason that most contributors to the Δ*G*_*membrane*_ and *G*_*protein*_ terms are unlikely to contribute to ΔΔ*G*_*exp*_ (Supplementary Note 1). In the following sections, we discuss how the *γ*Δ*A* term, specifically the interfacial tension in the membrane, and certain contributors to the Δ*G*_*membrane*_ term, specifically changes in K_A_ and k_c_, may explain our data in the context of existing models.

The term −*γ*Δ*A* describes the gain in energy from opening the channel, increasing the protein’s membrane area by Δ*A* with tension *γ*, and this is the core contribution to the *force-from-lipids* hypothesis; we assert this term best explains how the presence of polymer increases the channel activation pressure threshold. The applied tension acts against the lateral pressure profile induced by the membrane that surrounds MscL (Ollila et al., 2011). Therefore, lowering the water-bilayer interfacial tension leads to higher activation pressures (Melo et al., 2017). Mechanistically, MscL’s gating is driven by its tension-sensitive N-terminal helix which resides along the bilayer-water interface (Bavi et al., 2016), supporting our interpretation. Prior MD simulations suggest that MscL opening is most sensitive to changes in the head-group surface water-bilayer interfacial tension, leading to the approximation ΔΔ*G* = −Δ*γ*_*int*_Δ*A* (Melo et al., 2017; Ollila et al., 2011). Our simulations provide direct evidence for a lower interfacial tension in the membrane with applied tension due to recruitment of the PEO chains of the poloxamer polymers to the membrane-water interface (Figure 4K). However, simulations do not provide the magnitude of the reduction of interfacial tension. We can, however, benchmark the expected magnitude of the effect from air-lipid monolayers studies and compare these values to estimates derived from our measurements. Jordanova et al studied membranes containing the lipid DPPE with poloxamer P188 and found a change of interfacial tension of the lipid monolayer of about Δ*γ*_*int*_ ≈ −12 mN/m with adsorption of saturating polymer concentrations (Jordanova, Tenchov, and Lalchev 2011). We extract ΔΔ*G*_*exp*_ from the experimentally-measured changes in opening pressure Δ*P* for P124 at 0.1 % (Figure 2B) as 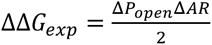, where *R* is the radius of the membrane patch in the pipette (*R* = 2.5 – 4 µm is a typical range in our experimental setup) (Wiggins and Phillips 2005). Assuming a Δ*A* of 20 nm^2^ (Chiang et al., 2004; Phillips et al., 2009), we find that ΔΔ*G*_*exp*_ and ΔΔ*G* are similar at 298K (2-3.2 x 10^-19^ Nm or 50-79 KbT and 2.3 x 10^-19^ Nm or 57 KbT, respectively). This concordance supports our proposed mechanism whereby P124-containing membranes increases MscL activation pressure thresholds primarily by reducing interfacial tension.

We next consider the key Δ*G*_*membrane*_ contribution between different membrane compositions (ΔΔ*G*_*membrane*_) in our study: the membrane-based, elastic costs of MscL opening. These elastic costs are proportional to the area expansion modulus K_A_ and bending rigidity k_c_ of the membrane (Wiggins and Phillips, 2005). We discuss other, likely insubstantial, contributions to ΔΔ*G*_*membrane*_ in Supplementary Note 1 (membrane asymmetry and hydrophobic mismatch). With respect to elastic membrane costs, Figures 2E, F show that the addition of P124 decreases K_A_ and k_c_, and following from published thermodynamic models, it appears that this should reduce the energetic cost of MscL opening and the activation pressure threshold of MscL (Δ*P*_*open*_) because the membrane should require less energy to deform to accommodate MscL’s open state (Nomura et al., 2012; Wiggins and Phillips, 2005). This is not the effect observed in Figure 2B and Figure 2-figure supplement 7A. To explain this apparent incongruity, we hypothesize that the addition of P124 to lipid membranes drives several, related elastic property changes which oppose one another. While polymer-containing membranes have lower K_A_ and k_c_ than lipid-only membranes (reducing the relative cost of opening a channel via ΔΔ*G*_*membrane*_), the polymer also lowers interfacial tension during membrane stretching (Figure 4K; reducing the thermodynamic driver of MscL opening, *γ*Δ*A*). To explain our results, we reason that the magnitude of ΔΔ*G*_*membrane*_ is smaller than the differences in *γ*Δ*A* between membrane compositions (50-79 KbT). This is consistent with order of magnitude estimates of Δ*G*_*membrane*_ contributors, the scaling of these contributors with K_A_ and k_c_, and our results from Figure 2E,F (Supplementary Note 1). Together, this suggests that although membrane elastic properties are important for MscL opening, and these properties correlate strongly with Δ*P*_*open*_, polymer-mediated effects on interfacial tension are likely the dominant mechanism driving MscL behavior in our study. When we focus on interfacial tension as the dominant property driving mechanosensitive channel activation, the relationships between K_A_ and k_c_ with activation pressure make sense. Because these properties are proportional to interfacial tension with *K*_*A*_ ∝ 4*γ*_*int*_ and *k_c_* ∝ *K*_*A*_*d*^2^, where d is the hydrophobic core thickness, changes in interfacial tension should proportionally alter these two membrane properties (Lipowsky and Sackmann, 1995), which is what we observed experimentally. In summary, we have compared our experimental results to available models for MscL gating and found that the polymer-induced changes of interfacial tension fit quantitatively to the *force-from-lipid* hypothesis. This property, which in turn alters the K_A_ and k_c_ values of membranes, is likely the dominant mechanism to explain the relationship between membrane properties measured here, membrane composition, and MscL gating.

Although most poloxamers we studied exhibited a monotonic effect on membrane properties and channel activation as a function of poloxamer concentration, we observed a non-monotonic effect of P184 concentration on MscLG22S activation and k_c_. While most poloxamers generally decreased K_A_ and k_c_ as a function of concentration (Figure 2, Figure 2-figure supplement 7, Figure 3-figure supplement 3), P184 induced a slight decrease in K_A_ at all concentrations while k_c_ decreased at low concentrations then returned to baseline levels at higher concentrations (Figure 3-figure supplement 3). Interestingly, we observed a similar effect between membrane k_c_ and MscLG22S activation pressure in the presence of P184 (Figure 3-figure supplement 2). We are unsure of the mechanism of this non-monotonic effect of P184 on k_c_ but attribute it to the large number of hydrophobic blocks relative to the small number of hydrophilic blocks in the polymer which likely leads to the polymer residing in a different location in the membrane relative to the other polymers explored (Rabbel et al., 2015). For example, P184 may increase membrane stiffness at higher concentrations through interdigitation (Battaglia and Ryan, 2005) or through changing the depth of P184 insertion in the membrane at high concentrations which is known to alter membrane properties (Grage et al., 2022; Zaki and Carbone, 2017). We speculate that these changes in polymer identity and integration at high concentration may also modulate polymer-mediated reduction in interfacial tension (as observed in Figure 4K for P124), for example by changing how the hydrophilic blocks adsorb to the bilayer under tension. In general, our results suggest that the effect membrane amphiphiles have on membrane properties can vary as a function of not only amphiphile identity, but also concentration, and may affect membrane mechanical properties in unexpected, nonmonotonic ways, further underscoring the importance of directly measuring membrane properties to determine the impact of membrane-associating biochemicals on physiological responses.

Finally, our observations of poloxamers interacting with membranes and altering mechanical properties and mechanosensitive channel activity highlight a potential unwanted effect of membrane amphiphiles on cellular membranes. Poloxamers represent a widely used class of nanoplastics. The concentrations of poloxamer we chose to assess in the present study are commonly used in laboratory cell culture applications to prevent cell sticking and improve cell integrity and are also found at these concentrations in household products (Guzniczak et al., 2018). Our findings suggest that poloxamers used at these concentrations may alter the behavior of membrane-embedded ion channels which in turn may affect environmental and human health or may affect biophysical studies in unexplored ways. In addition, these findings propose one potential mechanism for the alteration of biological function in response to nanoplastic pollution which has remained elusive (Fleury and Baulin, 2021)—modulation of global membrane mechanical properties. Our results suggest that nanoplastics which have been increasingly shown to interact with cellular membranes (Fleury and Baulin, 2021; Goodman et al., 2021; Hollóczki and Gehrke, 2019), may alter biological function by modulating membrane physical properties and the subsequent behavior of membrane-embedded proteins (Huang et al., 2022; Ohashi et al., 2020).

## Conclusion

In conclusion, we observed that various membrane-associating amphiphiles that decrease two membrane mechanical properties—membrane K_A_ and k_c_—increase the force required to activate mechanosensitive channels. Our results support a mechanism in which membrane interfacial tension, a modulator of both K_A_ and k_c_, regulates force transmission to mechanosensitive channels. Our study is the first to our knowledge to measure mechanosensitive channel activation as a function of membrane mechanical properties. By harnessing the unique features of polymeric amphiphiles, we could systematically alter several membrane properties by varying the amount of polymer present in the membrane. Our findings demonstrate that while K_A_ and k_c_ may be important for mechanosensitive channel function in some systems, they are not the dominating properties governing channel activation in our system containing non-natural amphiphiles. Rather, the relationships we observed between K_A_, k_c_, and channel activation pressure appeared to be a byproduct of alterations in a more dominant property, interfacial tension. These observations, using two distinct mechanosensitive channels, highlight a connection between the membrane physical environment and mechanosensitive channel function to show that bilayer properties, such as interfacial tension, are important mediators of mechanical force to channel proteins. Our results, accordingly, support the *force-from-lipids* mechanism of mechanosensitive channel activation by demonstrating that membrane composition and the associated bilayer mechanical properties can modulate mechanosensitive function through long-range, indirect interactions.

## Acknowledgements

This study was supported by the Air Force Office of Scientific Research YIP (FA9550-19-1-0039 P00001 to NPK) and the National Science Foundation (DMR-2145050 to NPK). This work was funded in part by the Chicago Biomedical Consortium with support from the Searle Funds at The Chicago Community Trust (to NPK and SMC). The authors acknowledge partial support from NIA/NINDS, National Institutes of Health (R01NS114413 to SMC), the Army Research Office (W911NF-22-2-0246), and the Department of Chemistry, College of Liberal Arts and Sciences, University of Illinois Chicago. MLJ was supported by Grant No. T32GM008382 from the National Institute of General Medical Sciences and the American Heart Association Predoctoral Fellowship under Grant No. 20PRE35180215. MJL was supported by NU’s Molecular Biophysics Training Program through NIH NIGMS (5T32 GM008382). PGD was supported by NIH NIDDK (1R56DK119709-01, 1R01DK123463-01). This work was supported by the Northwestern University Sanger Sequencing Facility. This research was supported in part through the computational resources and staff contributions provided for the Quest high performance computing facility at Northwestern University which is jointly supported by the Office of the Provost, the Office for Research, and Northwestern University Information Technology.

## Material and Methods

### Materials

Gibco G418 (geneticin), McCoy’s 5A medium, Opti-MEM transfection medium, Fetal Bovine Serum (FBS), CaCl2, MES sodium salt, BAPTA NA4, Poloxamer 188, 124, 407, and 184, Tris, B-casein, Fluo-4, MTSET, diphenylhexatriene, 12:0 LPC, molecular biology buffers and enzymes, and polyethylenimine 25K were obtained from Thermo Fisher Scientific. HEPES, NaCl, N-ethylmaleimide, and EGTA was purchased from Millipore Sigma. Human Osteosarcoma U-2 OS HTB-96 cell line was purchased from ATCC.

### Methods

#### Mammalian expression plasmid cloning

The *E.Coli* MscL-mEGFP fusion gene, EcMscLGFP, was amplified using polymerase chain reaction (PCR) and inserted into a mammalian pcDNA3.0 vector (Invitrogen) using restriction digest. A G22C mutation was introduced in the *mscl* gene using primers with a point mutation and the entire plasmid was amplified using PCR. The cloned plasmids were transformed into Top10 electrocompetent cells, midiprepped (PureLink Plasmid Midiprep Kit, Thermo Fisher), and plasmid inserts were confirmed using Sanger sequencing. TREK-1 was amplified using PCR and cloned into the pcDNA3.0 vector as a fusion to GFP by removal of EcMscL using restriction enzymes. Insertion and GFP fusion of TREK-1 was confirmed using Sanger sequencing. MscL and TREK-1 mutants were tagged with mEGFP on their C-terminus; see plasmid maps in Supplementary File 2. The MscL sequence was derived from pET19b:EcMscLGFP which was used previously as an *in vitro* membrane protein folding reporter (Addgene plasmid # 165097) (Jacobs et al., 2019). The TREK-1 sequence was from pGEMHE:mTREK-1(K271Q), which was a gift from Dan Minor (Addgene plasmid # 133270) (Lolicato et al., 2017).

#### Stable cell line transfection and selection

U2OS cells were plated in a 24-well plate and grown to 70% confluency on the day of transfection. Mammalian expression vectors containing EcMscLGFP or TREK-1GFP were transfected into U2OS cells using polyethylenimine (PEI). 500 ng DNA was mixed with 100 μL Opti-MEM transfection media and 2.5 μg PEI. This solution was incubated for 15 minutes at room temperature (20-25°C) then added dropwise to U2OS cells in 1 mL McCoy’s 5A media with 10% FBS. The transfection media was replaced with fresh McCoy’s 5A with 10% FBS and 300 μg/mL geneticin after overnight incubation. Transfected cells were selected using 300 μg/mL geneticin in McCoy’s 5A media with 10% FBS for 1 month or until the cells reached confluency in a T-75 75 cm^2^ flask. Cells were further propagated in McCoy’s 5A with 10% FBS and 100 μg/mL geneticin. Selection efficiency was confirmed using GFP fluorescence which was expressed as a fusion protein to the protein of interest.

#### GPMV formation and treatment with poloxamer

GPMV formation was performed using established techniques (Sezgin et al., 2012). Briefly, U2OS cells were plated at least one day prior to GPMV formation and grown to 70% confluency in a 6 well plate at the time of GPMV formation. GPMV buffer was used for all GPMV preparations and consisted of 10 mM HEPES, pH 7.4, 150 mM NaCl, and 2 mM CaCl2. N-ethylmaleimide (NEM) was prepared at 0.25 M in sterile water and stored in 300 μL aliquots. 30 μL NEM was added to 1 mL of GPMV buffer containing the indicated concentration of pre-dissolved poloxamer or detergent and mixed by vortexing. Cells were washed twice with GPMV buffer. The NEM-GPMV buffer solution was then added gently to the U2OS cells and incubated at 37 °C for 1 hour. GPMVs were observed and subsequently removed from adherent cells by gentle aspiration and were incubated for at least 3 additional hours at 4 °C prior to experiments to ensure poloxamer equilibration.

#### Fluorescence and phase-contrast microscopy

Cells and GPMVs were imaged using epifluorescence and phase-contrast microscopy to confirm transfection and protein transfer to GPMV membranes. Glass bottom, 96-well plates were passivated by incubating β-casein solution at 5 mg/mL in 10 mM Tris, pH 7.4 in each well for 30 minutes at room temperature. This solution was replaced with GPMV buffer at pH 7.4 and GPMVs were diluted to a concentration suitable to observe single GPMVs. GPMVs were imaged using differential interference contrast (DIC) and transfection efficiency and membrane protein integration were confirmed using epifluorescence with a GFP filter. All images being compared were brightness and contrast adjusted with identical settings and imaged at 20% light with identical exposure times. Images were taken on a Nikon Eclipse Ti2 Inverted Microscope, and analysis was performed using NIS software, ImageJ (Schindelin et al., 2012), and GraphPad Prism 9.

#### Micropipette aspiration

GPMV membrane area expansion modulus (K_A_) and bending rigidity (k_c_) were measured using micropipette aspiration techniques (Evans et al., 1976). Micropipettes were constructed from borosilicate glass capillaries (World Precision Instruments) pulled using a P-1000 micropipette puller (Sutter instruments). The end was blunted and polished to an inner diameter of 5–8 *μ*m using a microforge. The micropipette and sample chamber surfaces were passivated using 5 mg/mL β-casein for 1 hour and were filled with GPMV buffer, pH 7.4. 10 μL GPMVs prepared in the presence of poloxamer, detergent, or no additive were diluted in a 1 mL glass chamber in GPMV buffer, pH 7.4 (10 mM HEPES, 150 mM NaCl, 2 mM CaCl2). Imaging was performed on a Nikon inverted microscope connected to a Validyne pressure sensor and digital manometer (model DP 15-32; Validyne Engineering, Northridge, CA). A Narishige micromanipulator was used to control the micropipette (model WR-6; Narishige). GPMVs free of visible defects were aspirated and the membrane areal response to a change in pressure was measured using a series of DIC images with pressure recordings. Membrane K_app_ is calculated from the slope of the percent area dilation within the pipette (ΔA/Ao) and membrane tension (τ),

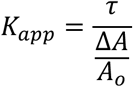

where ΔA is the change in area of GPMV membrane within the micropipette and Ao is the initial area of the aspirated GPMV within the pipette. Compared to previous studies K_app_ was measured at elevated tensions that recruit any membrane area reservoirs and thus probe the elastic response of the membrane (Steinkühler et al., 2021). Bending elastic modulus, k_c_ is calculated from the same aspiration measurements through the following equation in the low-tension regime (<0.5 mN/m),

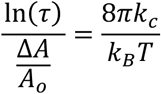

where kB is the Boltzmann constant, and T is temperature. Membrane K_A_ is calculated through correction of K_app_ with k_c_ by subtraction for relaxation of thermal undulations measured by k_c_ (Rawicz et al., 2000; Zhou and Raphael, 2005). For each stress/strain measurement in the area expansion regime, the contributions of thermal undulations (Δα(i)) was subtracted from the areal strain (ΔA/Ao),

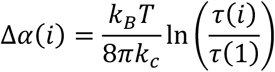

where τ(1) is the initial tension. The corrected areal change was then plotted against τ and the slope of this plot was used to calculate K_A_.

### Patch Clamp electrophysiology of stretch-activated ion channels

Channel activity measurements were performed in identical conditions as micropipette aspiration. GPMVs containing EcMscL and TREK-1 prepared under indicated poloxamer conditions were added to an open-air bath on an Olympus IX73 phase-contrast microscope. Electrophysiology pipettes were pulled using a Sutter P-1000 micropipette puller and fire polished using a microforge to 5-8 MΩ resistance. Voltage-clamp measurements were recorded on a Molecular Devices Axon Instruments system including an amplifier and digitizer Low-Noise Data Acquisition System plus Hum silencer along with an ALA Scientific Instruments High Speed Pressure Clamp pressure controller which was used to apply a pressure ramp during electrophysiology recordings. All signals were integrated using Clampex software and pressure and current values were recorded temporally. A pressure ramp from 0 mmHg to -160 mmHg was applied at a constant rate for all recordings. EcMscL and TREK-1 recordings were performed at -10 mV and -40 mV, respectively unless channel conductance calculations were being measured. Pressure and current were recorded through these experiments and were used to calculate pressure sensitivity and channel conductance, respectively. Each pressure sensitivity recording was performed on an independent GPMV and pressure ramps were never repeated as a decrease in pressure sensitivity is observed after repeat pressure ramps because of an increase in membrane tension when the membrane patch diffuses up the pipette (Nomura et al., 2012). All electrophysiology recordings were performed at 25 °C. Bath solution was composed of 10 mM HEPES, pH 7.4, 150 mM NaCl, 2 mM CaCl2 for MscL and 10 mM HEPES, pH 7.4, 150 mM KCl, 2 mM CaCl_2_ for TREK-1. The pipette solution was composed of 10 mM HEPES, pH 7.4, 100 mM NaMES, 10 mM BAPTA Na4, 10 mM NaCl, 10 mM EGTA. All recordings were performed in the cell-attached or excised-patch configuration. A seal > 8 GΩ was used as criteria for analysis of single-channel observations.

### Electrophysiology data analysis

Electrophysiology experiments were quantified using Clampfit software which displayed the current and pressure during the voltage-clamp recording. EcMscL and TREK-1 channel activation sensitivities were quantified using the pressure at first channel opening which has been described previously (Nomura et al., 2012). Many recordings only contained one channel, and this first channel was clearly distinguished from electrical noise. However, some recordings contained many channels which could not be distinguished from membrane unsealing, making maximum current an unreliable measurement. Membrane rupture pressure was calculated by the pressure at which a very large current change occurred, indicating complete membrane rupture. Single channel conductance was measured by calculating the current through single channels at various voltages and was independent from pipette pressure. Each current and voltage measurement was recorded from an independent GPMV, and this measurement was not repeated by any single GPMV to mimic pressure sensitivity measurement conditions. The slope of many of these I/V relationships was used to calculate channel conductance. TREK-1 was observed to behave similarly to previous studies and had a short dwell time and was potassium selective with a much smaller conductance than MscL (Xian Tao Li et al., 2006).

### Mass Spectrometry

Mass spectrometry (MS) was used to characterize the integration of poloxamer into the GPMVs. Samples were prepared as described for GPMV formation and free poloxamer was removed by centrifugation at 12,000 x *g* for 30 minutes. Pelleted GPMVs were stored at -80 °C until analysis. Matrix-assisted laser desorption/ionization time of flight (MALDI-TOF) MS was carried out using an Autoflex® speed MALDI-TOF (Bruker Daltonik GmbH,Bremen Germany). A 10 mg/mL matrix solution of α-cyano-4-hydroxycinnamic acid (CHCA) in 70% ACN in water with 0.1% trifluoroacetic acid (TFA) was prepared. Using the dried drop (DD) method, aqueous solutions of P124-only, P188-only standards and P124-GPMV, P188-GPMV integrated samples were mixed with the matrix at a ratio of 1:1 (vol/vol). For each sample, 5000 shots were fired using the “random walk” function. Positive ion data was collected in both the linear and reflectron modes with external calibration. The acquisition mass ranges were *m/z* 1000–4000 and *m/z* 4000–12500 for P124 samples and P188 samples, respectively. For the poloxamer(P)-GPMV samples, mass range for the P-GPMV lipids was *m/z* 500–1000.

### Anisotropy

Fluorescence anisotropy measurements were used to determine relative membrane fluidity as described previously (Jacobs et al., 2021). 1 mL stock GPMVs were prepared and unbound poloxamer micelles were removed from solution through centrifugation at 10,000 x *g* for 1 minute, removing supernatant solution, and resuspension of GPMVs in 1 mL GPMV buffer. A diphenylhexatriene (DPH) stock solution was prepared by adding dry DPH to GPMV buffer at 100 μM. This solution was mixed 1:1 with purified GPMVs and anisotropy was measured using an Agilent Technologies Cary Eclipse Fluorescence Spectrophotometer with an Automated Polarization Accessory. Fluorescence anisotropy was assessed at excitation 360 nm and emission 435 nm and was calculated by software provided with the instrument,

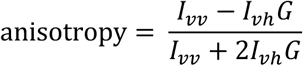

where Ivv and Ivh are the intensities of the vertically and horizontally polarized light, respectively, after excitation with vertically polarized light. G = Ihv/Ihh which is a grating correction factor for the optical system.

### Chemical activation of MscL and calcium imaging

Fluo4, a calcium indicator which has been used previously to sense mechanosensitive channel behavior (Cahalan et al., 2015), was used to determine if P124 inhibits MscL activation through inhibition of pore hydration. Briefly, Fluo4 was mixed with GPMV buffer to a 1x dye concentration. GPMV formation solution was prepared by adding 30 μL of 0.25 M NEM to 1 mL of diluted Fluo4 solution in GPMV buffer. Cells were washed twice with GPMV buffer and incubated in GPMV formation solution for 1 hour at 37 °C. Free-floating GPMVs were isolated by transferring formation solution to an Eppendorf tube and incubated for 3 additional hours. A glass-bottom 96-well plate was passivated by incubating with a β-casein solution in 10 mM Tris, pH 7.4 for 30 min at room temperature. The wells were rinsed 3 times with GPMV buffer and 100 μL GPMV buffer with 5 mM CaCl2 was added to each desired well. 10 μL GPMVs were added to the bottom of each well. 5 μL of 21 mM MTSET was added to each well and GPMVs were incubated for 30 min at room temperature. Ca^2+^ influx through MscL opening was quantified from raw images by a fluorescence increase in the GFP channel for each GPMV visible in the DIC channel of a Nikon Ti2 Inverted Microscope.

### Molecular Dynamics Simulations

P124 polymer was assembled using CHARMM-GUI Polymer Builder from PEO and PPO groups from the CHARMM36 forcefield (Choi et al., 2021; Jo et al., 2008). Based on previous work (Rabbel et al., 2015) we assumed that P124 inserts as a harpin into the membrane. We further assumed that the incubation time is sufficient for P124 to flip-flop in the membrane and assume a symmetric configuration. The polymer chains were pre-equilibrated in water and a harpin-like conformation was selected by hand. Varying number of polymer chain copies were then inserted to a pre-equilibrated DOPC membrane (300 lipids in total) as indicated by the mol fraction reported in Figure 4H,I and Figure 4-figure supplement 3. The parallel periodic membrane images were separated by a 4 nm water layer. Simulations were run using GROMACS (2020) with a timestep of 2 fs, Verlet cutoff-scheme, electrostatics and LJ interactions cutoff at 1.2 nm and PME long-range electrostatics. Simulations were run NPT ensemble, using a Nose-Hoover thermostat and semi-isotropic Parrinello-Rahman pressure coupling for a (laterally) tensionless membrane. Temperature coupling was done with the Nose-Hoover algorithm. The membrane elastic modulus *K*_*A*_ was estimated from the thermal bilayer lateral area fluctuations from a (fully equilibrated) 500 ns long trajectory (Feller and Pastor, 1999). Membrane thickness was calculated from the lipid headgroup phosphate distances between the two leaflets using MDAnalysis version 0.20.1 (Michaud-Agrawal et al., 2011). Statistical uncertainties in the membrane elastic modulus and thickness were estimated using blocking analysis (Flyvbjerg and Petersen, 1989) implemented in pyblock (James Spencer, http://github.com/jsspencer/pyblock) with block sizes chosen to remove temporal correlation from the data set (Wolff, 2004). To induce, membrane tension in the experiment for Figure 4K, the simulation box size was fixed to induce a positive area strain. Specifically, we considered the simulation of a single P124 harpin per membrane leaflet that in the membrane plane had an average area of 10.2 x 10.2 nm^2^ at zero membrane tension. Fixing this area to 10.8 x 10.8 nm^2^ induced an area strain of about 12% and included a mechanical tension of 17 mN/m (calculated using gmx energy, GROMACS).

## Supplemental Figures

**Figure 1–figure supplement 1.**
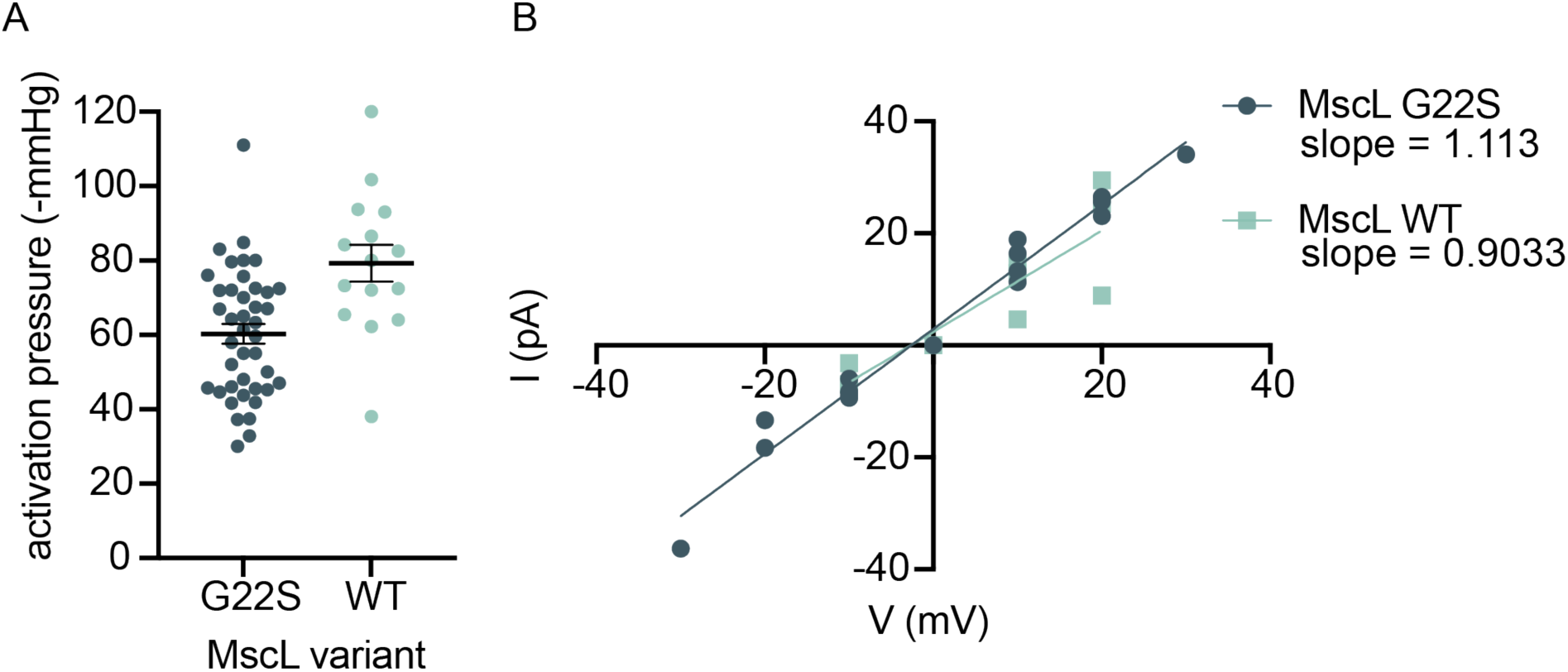
MscLG22S reduces the pressure required for activation compared to MscLWT but does not change channel conductance. A) MscL activation pressure is reduced in the G22S mutant. The pressure at first channel opening was measured to quantify MscL activation sensitivity. Individual points represent independent GPMV measurements. Mean is plotted with error bars which represent standard error of the mean. B) Conductance calculations for MScLG22S and MscLWT demonstrate that MscL channel size and selectivity is not affected by the G22S mutation. MscLG22S conductance = 1.113 nS (95% confidence interval 1.020 – 1.207). MscLWT conductance = 0.9033 nS (95% confidence interval 0.5761 – 1.231)

**Figure 1–figure supplement 2.**
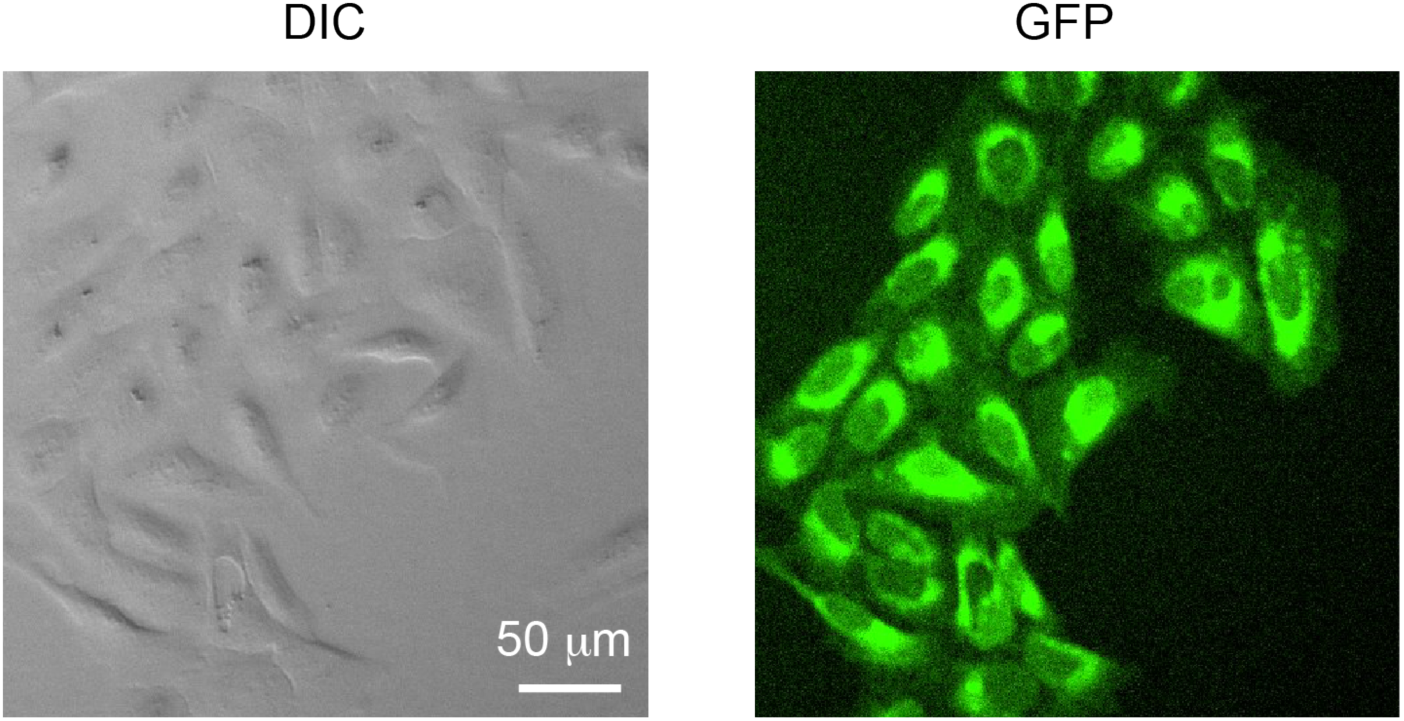
U2OS cell lines stably expressing MscLG22S tagged with GFP. MscLG22S expressed as a fusion protein with GFP was transfected into U2OS cells and selected using geneticin for 4 weeks. MscL is localized to various membranes within the cell including the plasma membrane.

**Figure 1–figure supplement 3.**
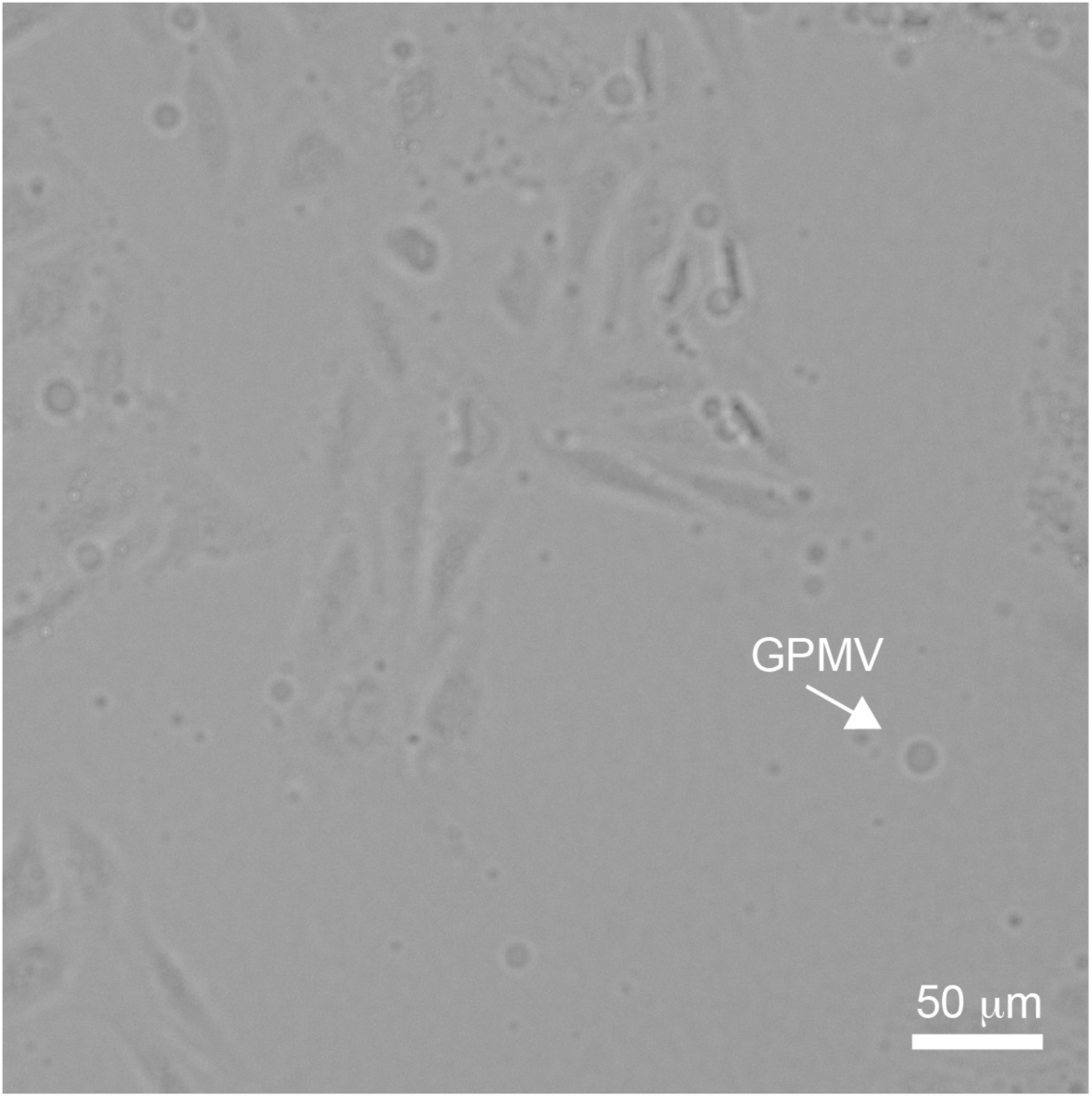
GPMVs budding off U2OS cells. DIC micrograph demonstrates Giant Plasma Membrane Vesicles (GPMVs) formation by budding off from adherent mammalian cells. This process creates large diameter, bilayer vesicles comprised of the cellular plasma membrane without cytoskeletal attachments or intracellular membranes.

**Figure 1–figure supplement 4.**
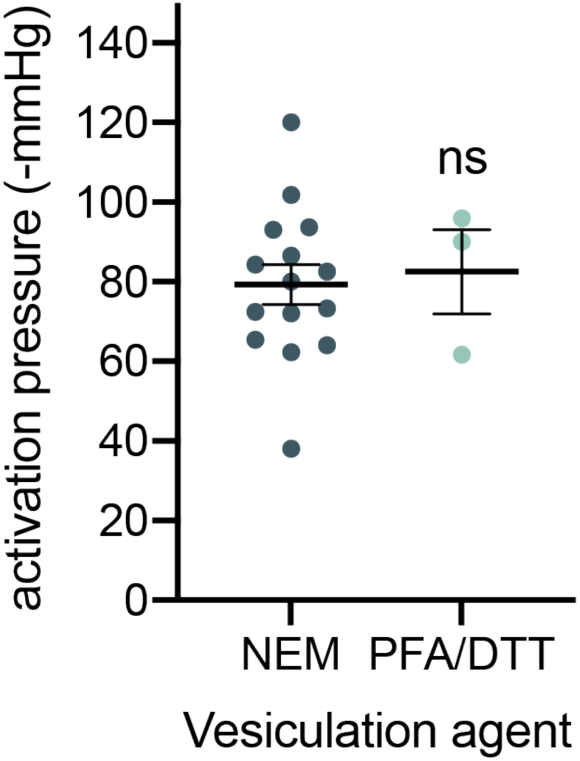
Differing GPMV formation methods yield similar MscL activation pressures, but crosslinking reagents may reduce the probability of channel opening. GPMV formation is commonly performed using two different methods, NEM or PFA/DTT. While PFA/DTT is the most commonly used reagent mixture, this method induces crosslinking of lipids and proteins that is not amenable to mechanosensitive channel activation. In the presence of PFA/DTT, after >30 attempts over three independent GPMVs preparations, only three instances of functional MscL channels were observed. In contrast, NEM preparation allowed reliable observation of MscL activation. The two methods for GPMV formation did not demonstrate different activation pressure thresholds of MscL. p-values to determine significance between means was measured using Student’s t-test. non-significant (ns) p > 0.05, n > 30.

**Figure 1–figure supplement 5.**
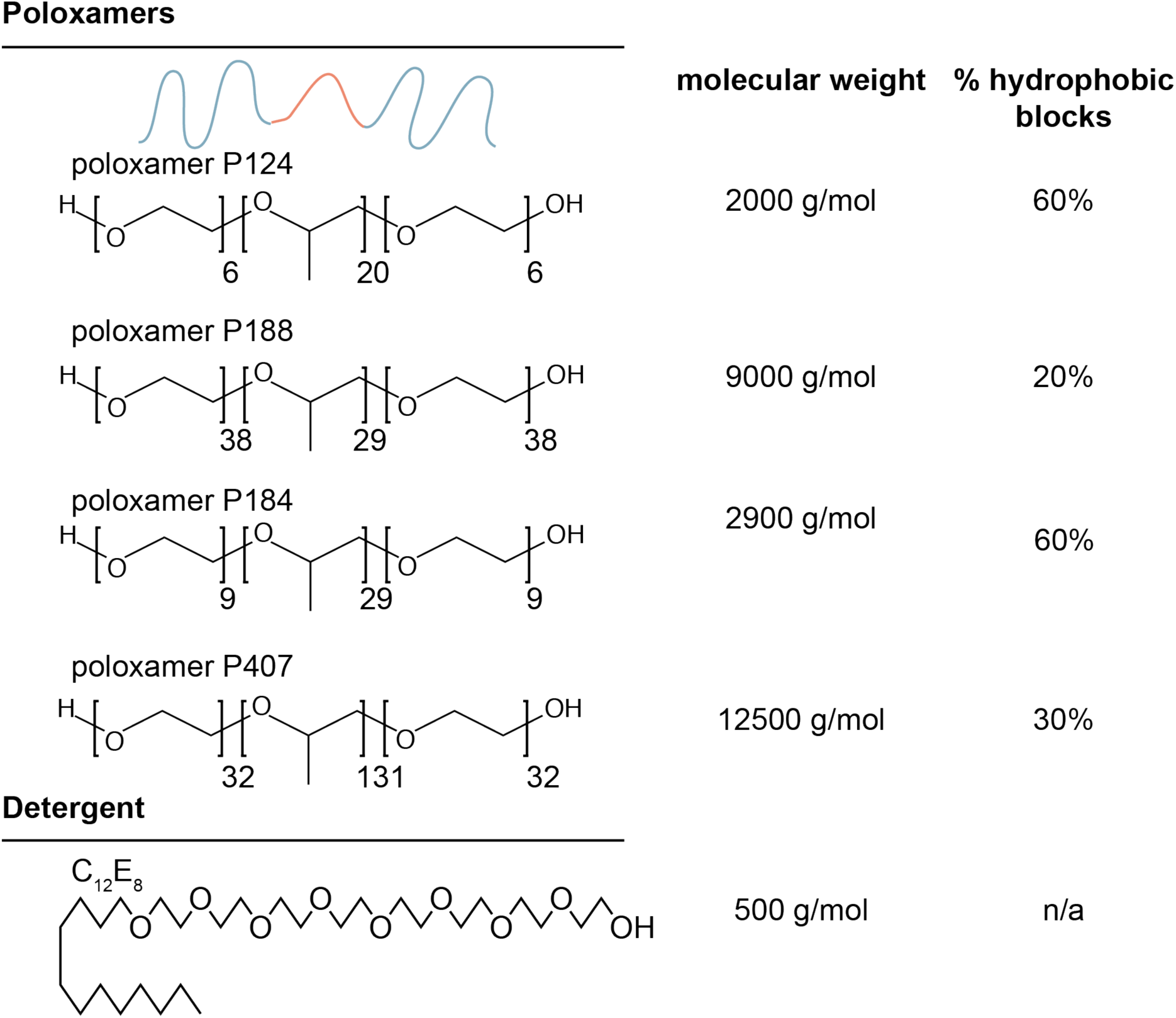
Schematic and properties of amphiphiles used in this study. We used various synthetic amphiphiles to exogenously alter membrane mechanical properties. These amphiphiles belong to two classes, poloxamer (Pluronic) and small nonionic detergent. Poloxamer is a class of triblock copolymers which consist of a polypropylene oxide hydrophobic block sandwiched by two polyethylene oxide hydrophilic blocks. Poloxamers have similar properties to surfactants in their ability to insert into bilayers but have much larger molecular weights. The detergent C12E8 we used has a much smaller molecular size in comparison to poloxamer.

**Figure 1–figure supplement 6.**
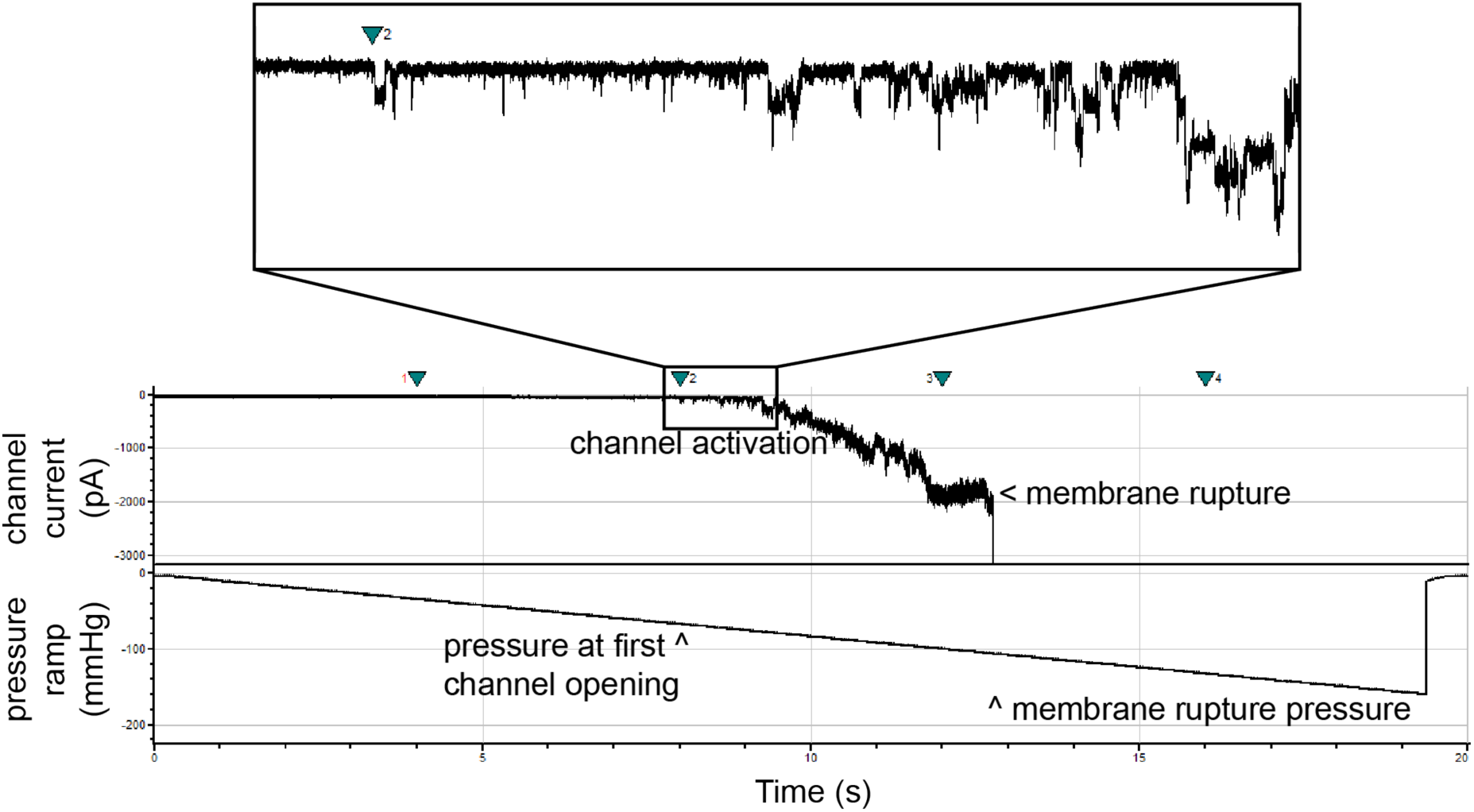
Sample electrophysiology data analysis workflow. We calculated MscLG22S pressure sensitivity using an integrated pressure controller to quantify the pressure at which the first MscLG22S channel activated. Higher pressures open more than one channel as demonstrated by multiples of original channel current. Inset demonstrates stochastic channel openings due to the applied pressure. We applied enough pressure to pop the GPMV membrane to ensure the pressure was high enough to activate mechanosensitive channels in the GPMVs and to enable the quantification of membrane stability.

**Figure 1–figure supplement 7.**
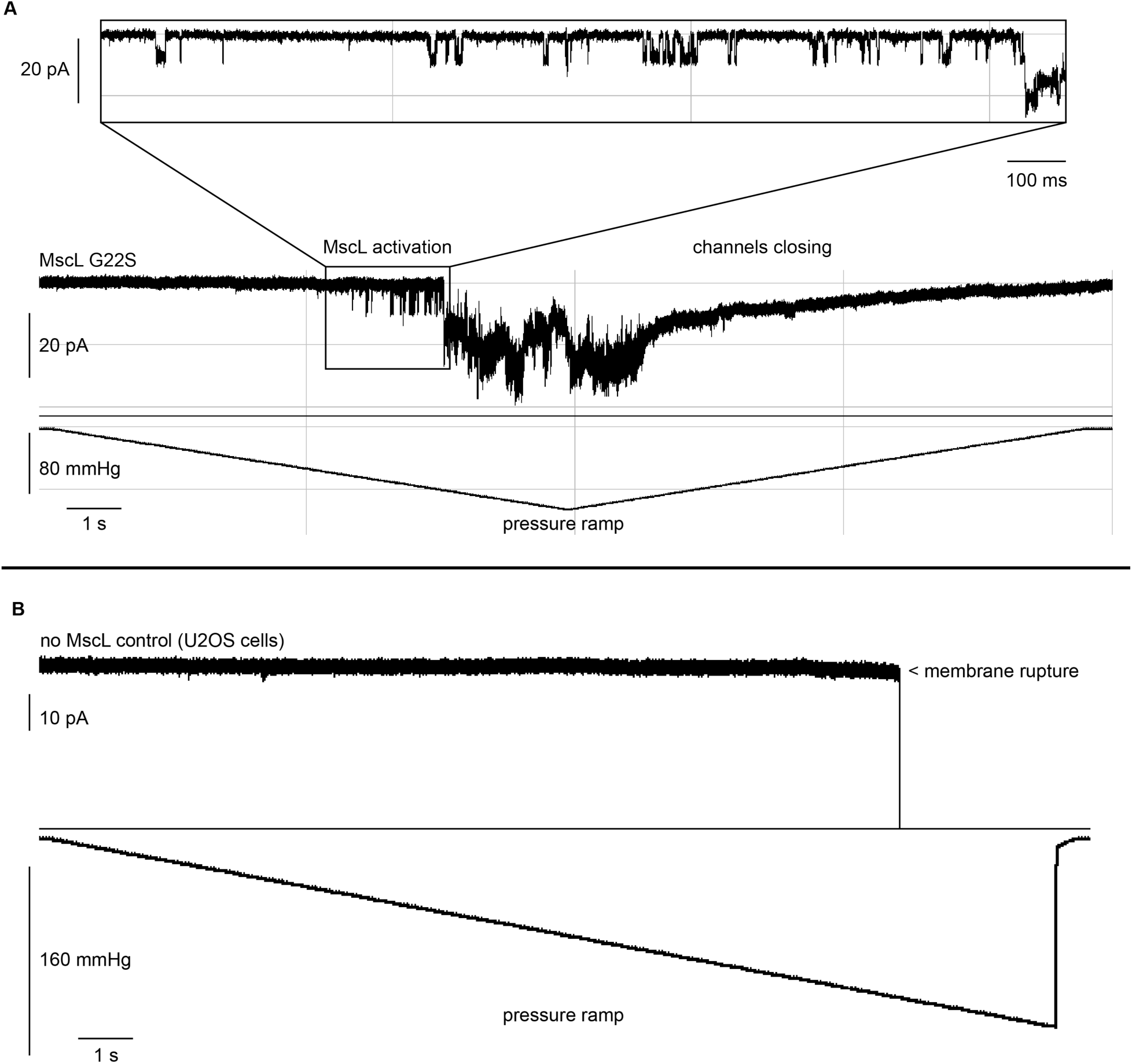
Example results of full traces of MscLG22S and no MscL electrophysiology recordings. A) MscLG22S exhibits stochastic activation behavior in response to pressure in GPMVs. When applied pressure is returned to 0 mmHg, MscLG22S closes and current returns to baseline. This observation suggests that the change in current is due to channel activation and not membrane rupture. B) No MscL control demonstrates no channel currents under MscLG22S activation conditions. Channel currents are not observed up to pressures high enough to rupture the membrane. The open/close pressure ramp in (A) was only used to demonstrate channel closure after pressure reduction. All other recordings were performed using the rate of pressure change in (B) for all MscLG22S activation calculations.

**Figure 1–figure supplement 8.**
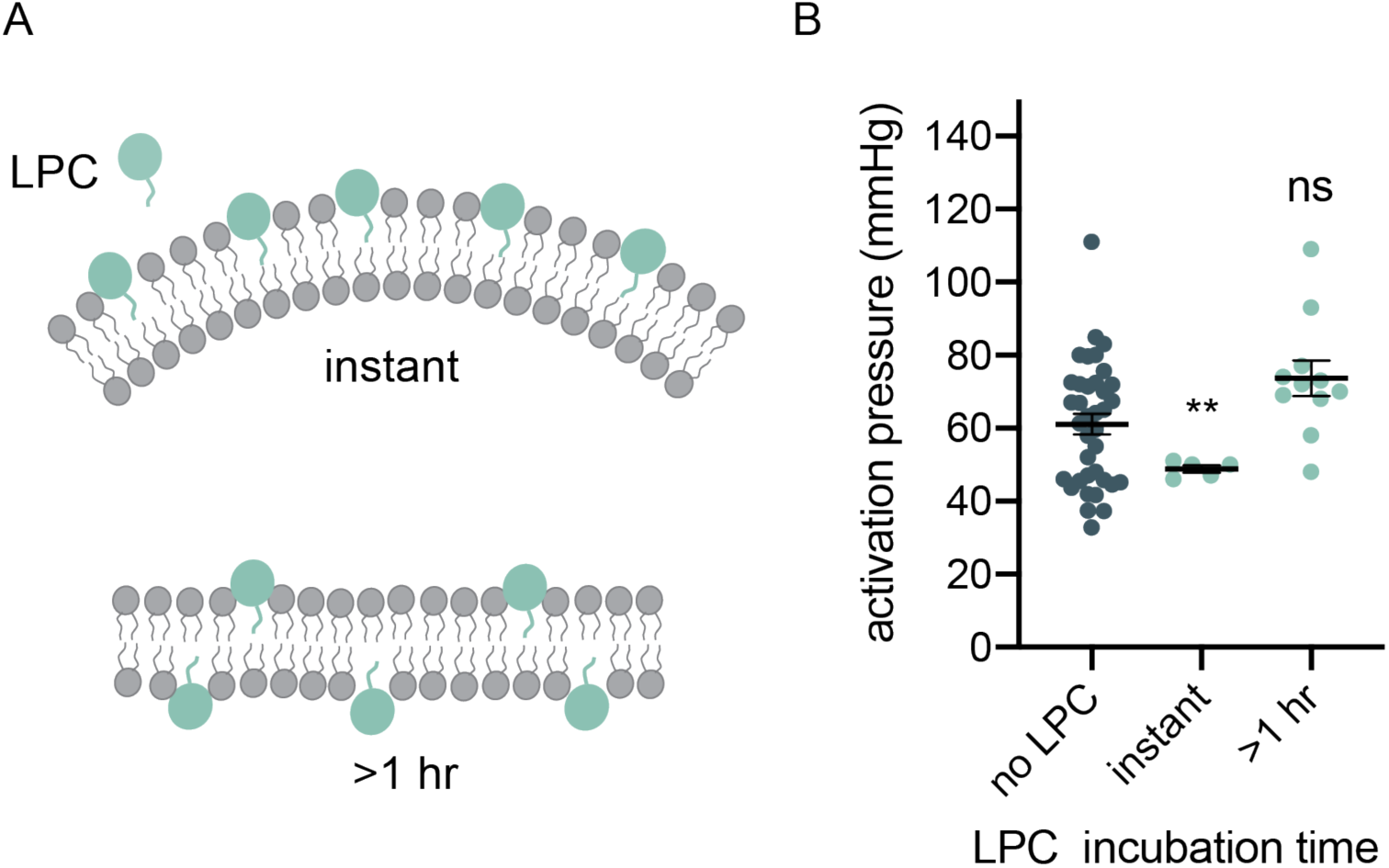
Lysophosphatidylcholine (LPC) activation confirms MscLG22S behavior in GPMVs formed from U2OS cells. A) LPC inserts into membranes and induces local changes in curvature on short timescales. These changes in curvature are known to reduce the pressure required to activate MscL (Nomura et al., 2012). On longer timescales, LPC equilibrates across bilayer leaflets, and this equilibration neutralizes curvature such that there is no expected effect on MscLG22S activation. B) MscLG22S activation pressure is reduced upon LPC introduction. After incubation for >1hr, LPC no longer alters MscLG22S activation pressure as expected. p-values were generated by ANOVA using Dunnett’s multiple comparisons test compared to no poloxamer. ** p ≤ 0.01, non-significant (ns) p > 0.05, n> 10.

**Figure 2–figure supplement 1.**
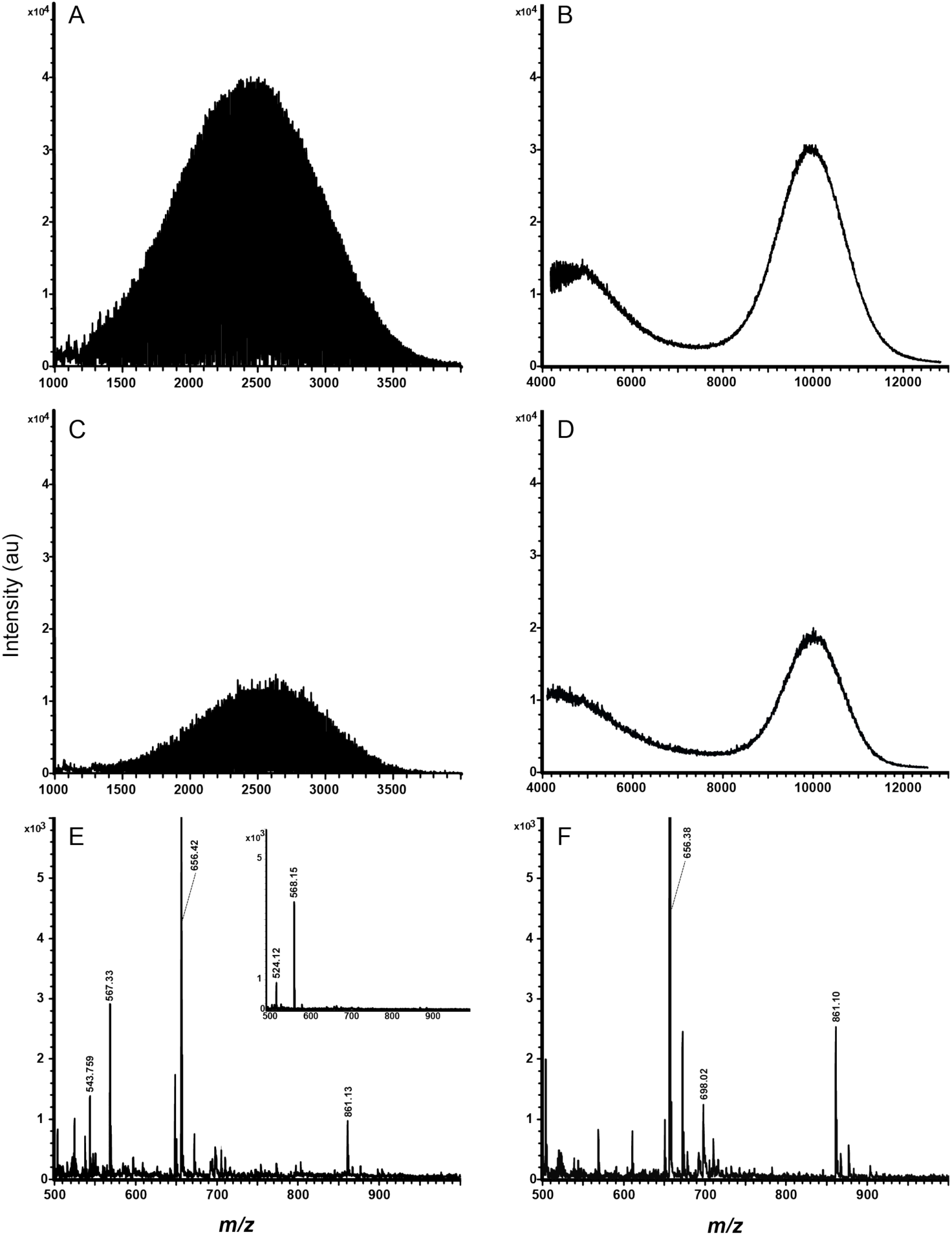
MALDI-TOF mass spectra of pure poloxamer A) P124 and B) P188. Samples in panes C–E are poloxamer-integrated GPMVs. In separate experiments, data was acquired in the higher mass regions, *m/z* 1000–4000 and *m/z* 4000–12500 to detect C) P124 and D) P188 in GPMVs, respectively. Also, data was acquired in a lower mass region *m/z* 500–1000 to detect E) GPMV lipids of the P124-GPMVs and F) GPMV lipids of the P188-GPMVs. The inset in pane E is the matrix, CHCA, collected in the mass range *m/z* 500–1000.

**Figure 2–figure supplement 2.**
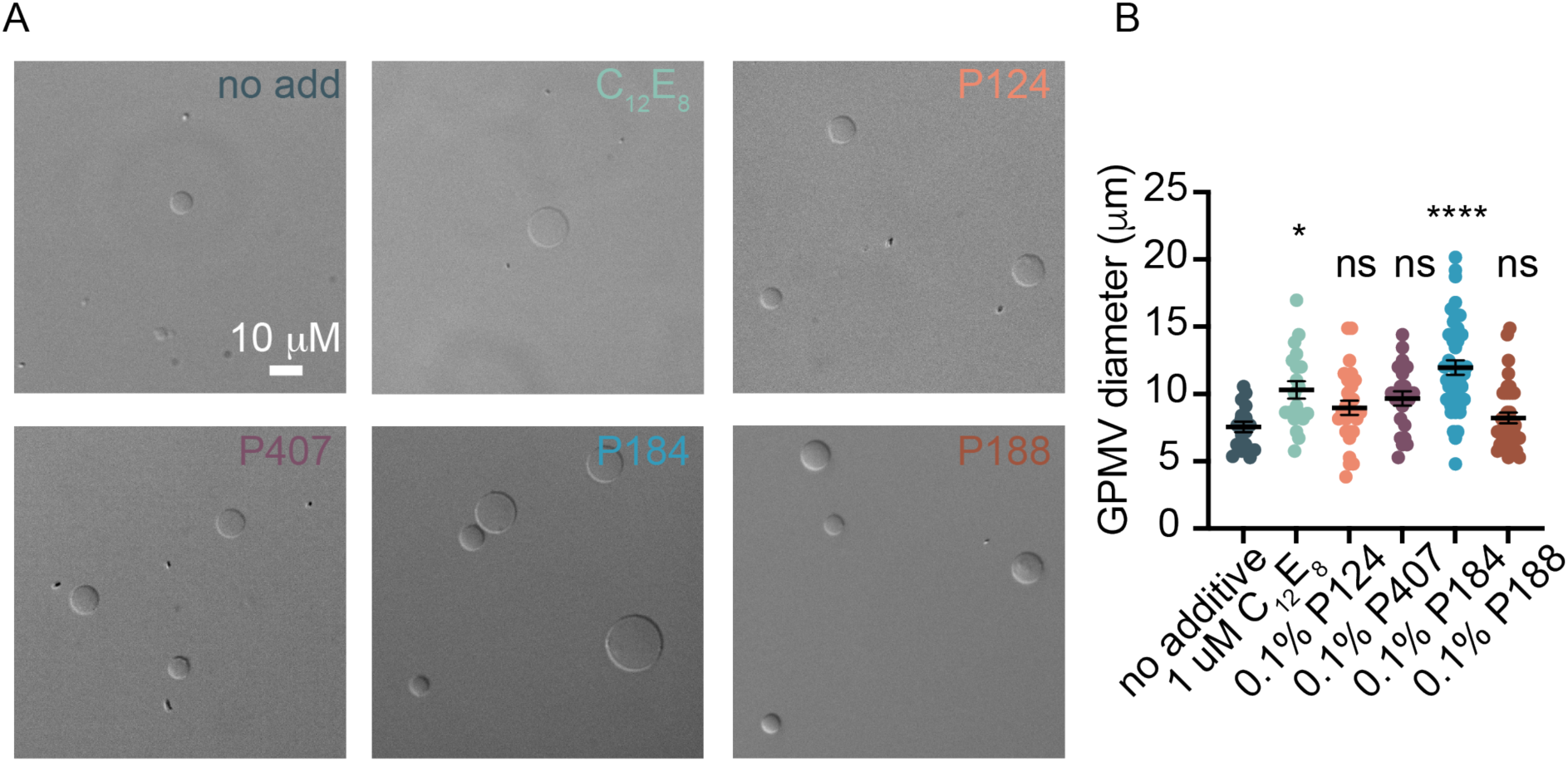
GPMV diameter increases in the presence of various poloxamers and a detergent. A) DIC phase contrast micrographs demonstrate the effect poloxamer and detergent (C12E8) on GPMV size. B) GPMV diameter was quantified for all GPMVs large enough to quantify. Individual points represent the diameter of a single GPMV, mean and standard error of the mean are plotted. p-values were generated using ANOVA Dunnett’s test of multiple comparisons compared to no poloxamer. **** p ≤ 0.0001, * p ≤ 0.05, non-significant (ns) p >0.05, n > 10.

**Figure 2–figure supplement 3.**
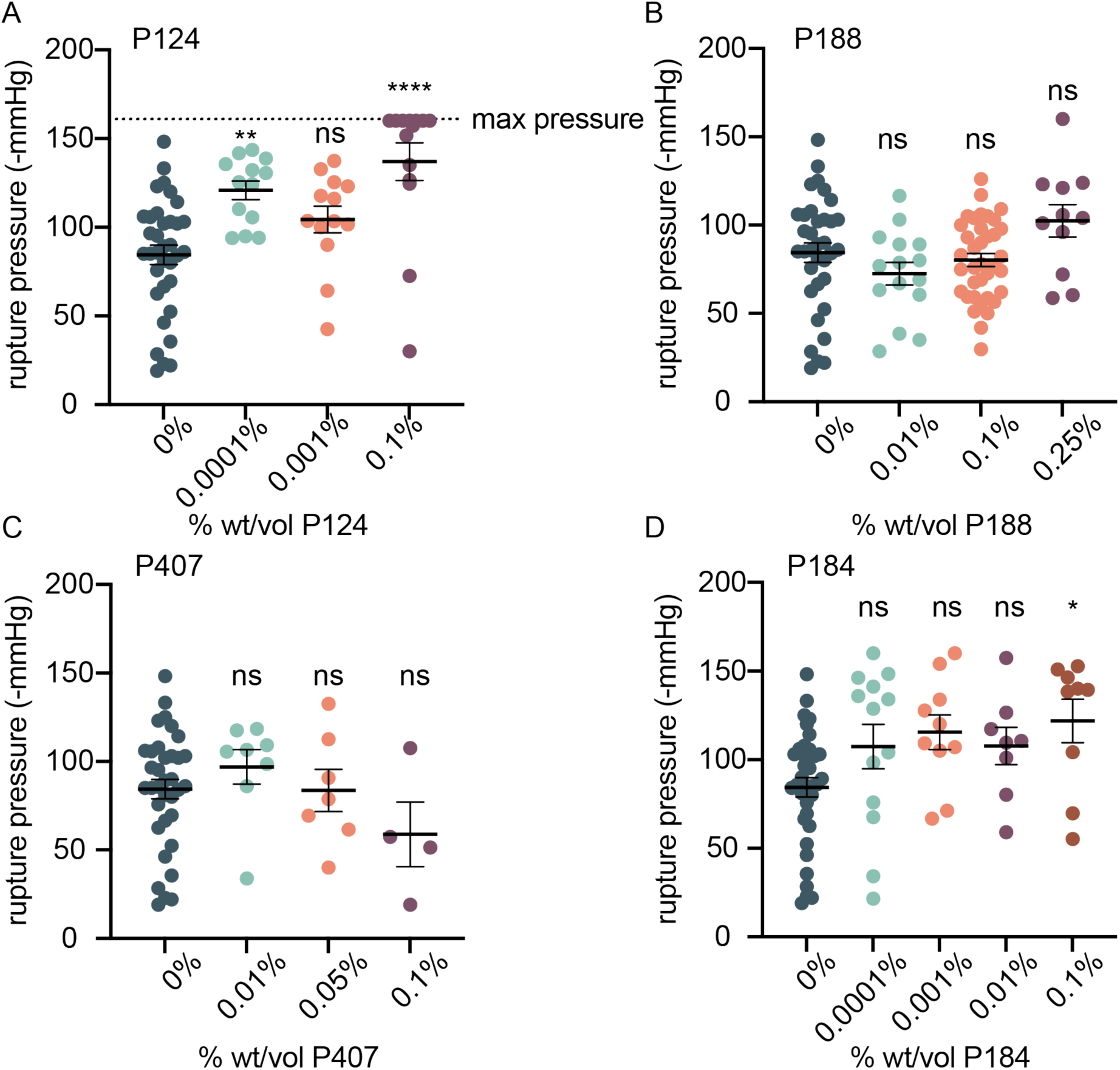
Poloxamer effect on membrane stability. We quantified membrane rupture pressure during our electrophysiology recordings of MscLG22S activation to measure if various poloxamers exhibited a strengthening effect on the membrane as has been previously discussed (Zaki and Carbone, 2017). The effect of P124 (A), P188 (B), P407 (C), and P184 (D) were measured, and we observed a membrane strengthening effect in the presence of some poloxamers. However, P407 exhibited a decrease in membrane stability which we hypothesize is due to the large molecular weight of this poloxamer. Each point represents the rupture pressure of a single GPMV, mean is plotted, and error bars represent standard error of the mean. p-values were generated by ANOVA using Dunnett’s test for multiple comparisons compared to no poloxamer (0%). * p ≤ 0.05, non-significant (ns) p > 0.05, n >10 for most samples except P407 (n > 3) which was prone to popping prior to the initiation of the electrophysiology recording.

**Figure 2–figure supplement 4.**
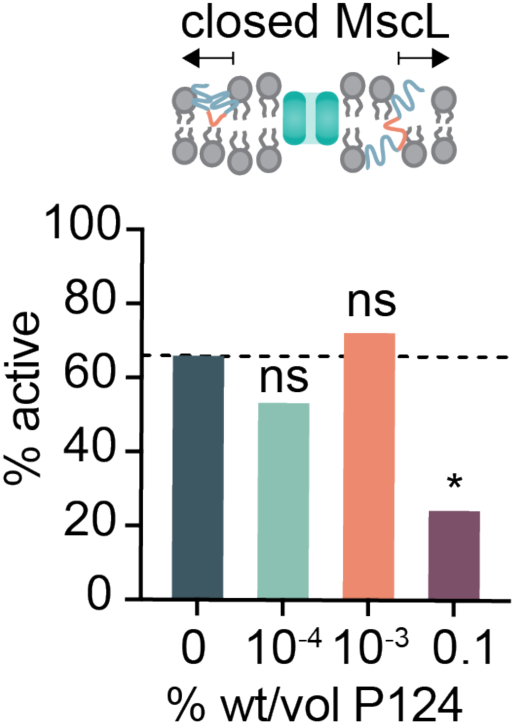
P124 prevents MscLG22S activation at high concentrations. Some electrophysiology recordings contained membranes that remained stable in the presence of high pressure (> -80 mmHg), yet MscLG22S did not activate. Recordings in which we observed activation of MscLG22S are presented as a percent of all recordings for a given condition (% active). The dashed line represents the percent of recordings in which MscLG22S openings were observed in the absence of poloxamer. n>12 for all conditions. p-values were generated using Fisher’s exact test comparing the distribution of values to 0% P124, * p ≤ 0.05, non-significant (ns) p >0.05, n>10.

**Figure 2–figure supplement 5.**
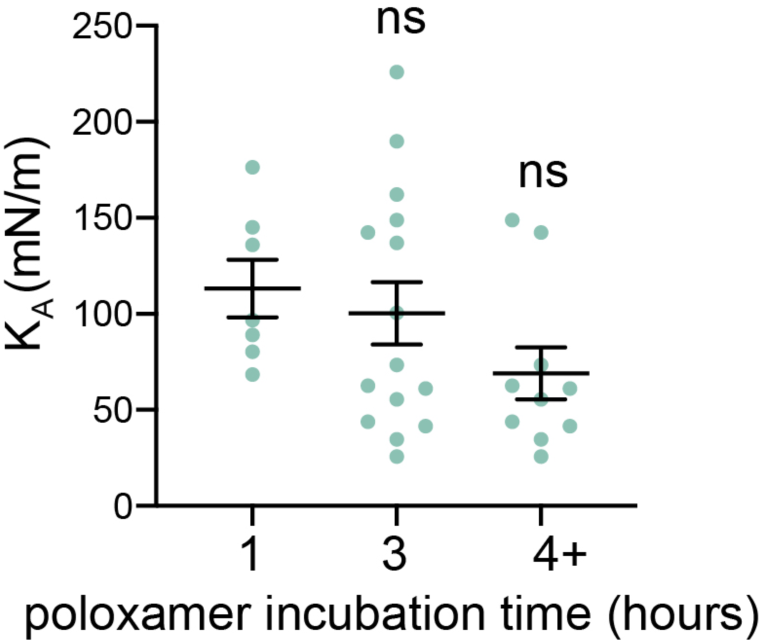
Poloxamer incubation time alters poloxamer effect on membrane properties. Incubation time noticeably, but does not significantly, alter membrane K_A_ in this experiment. GPMVs were incubated with 0.1% wt/vol P188 at 4 °C for either 1, 3, or 4 hours and membrane K_A_ was measured using micropipette aspiration techniques. p-values were generated by ANOVA using Dunnett’s test for multiple comparisons compared to 1-hour, non-significant (ns) p > 0.05, n > 6 independent GPMVs.

**Figure 2–figure supplement 6.**
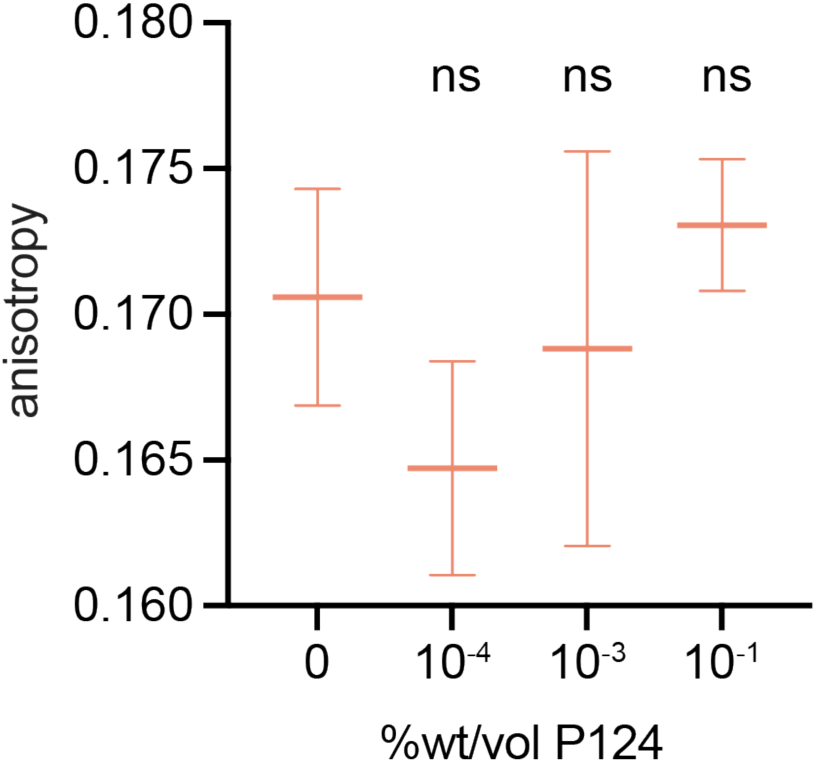
P124 effect on membrane fluidity. Fluorescence anisotropy was used to measure relative differences in membrane fluidity between GPMV membranes containing various amounts of P124. Decreased anisotropy indicates a relative increase in membrane fluidity. Mean and standard error of the mean are plotted. p-values were generated by ANOVA using Dunnett’s multiple comparisons test compared to no poloxamer, non-significant (ns) p >0.05, n>10.

**Figure 2–figure supplement 7.**
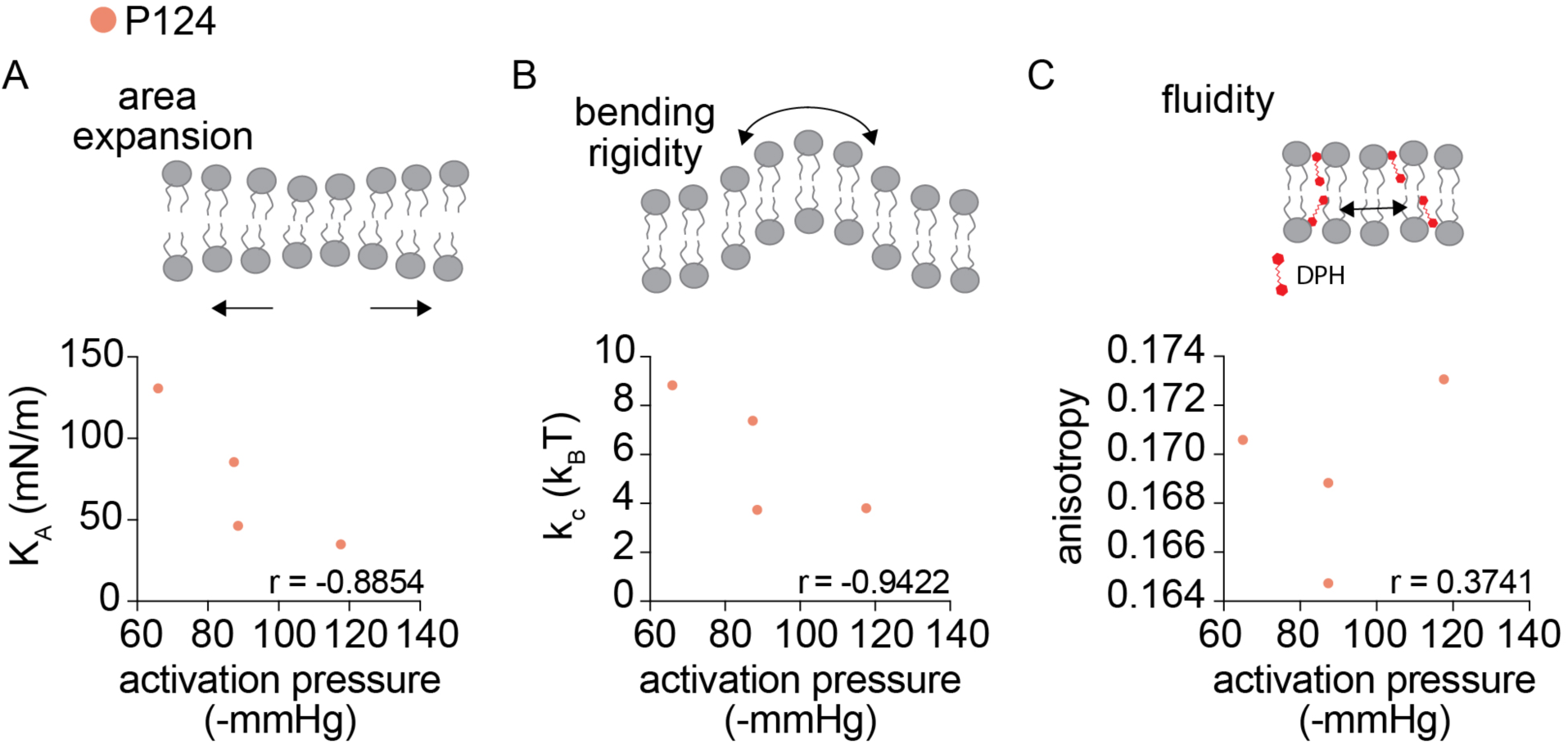
Poloxamer P124 alters GPMV mechanical properties. A) P124 decreases K_A_ and increases MscLG22S activation pressure. K_A_ is plotted as a function of MscL activation pressure. B) P124 decreases k_c_ and increases MscLG22S activation pressure. Mean k_c_ is plotted as a function of mean MscLG22S activation pressure. C) Membrane fluidity does not correlate with MscLG22S activation pressure. Membrane anisotropy was used to measure relative changes in membrane fluidity in the presence of P124. Decreased anisotropy indicates a relative increase in membrane fluidity. GPMVs were mixed with diphenylhexatriene (DPH) to a final concentration of 50 μM and fluorescence anisotropy was measured using a fluorimeter with fluorescence polarization capabilities. For all plots, Pearson’s r was calculated to measure correlation strength, n>10.

**Figure 3–figure supplement 1.**
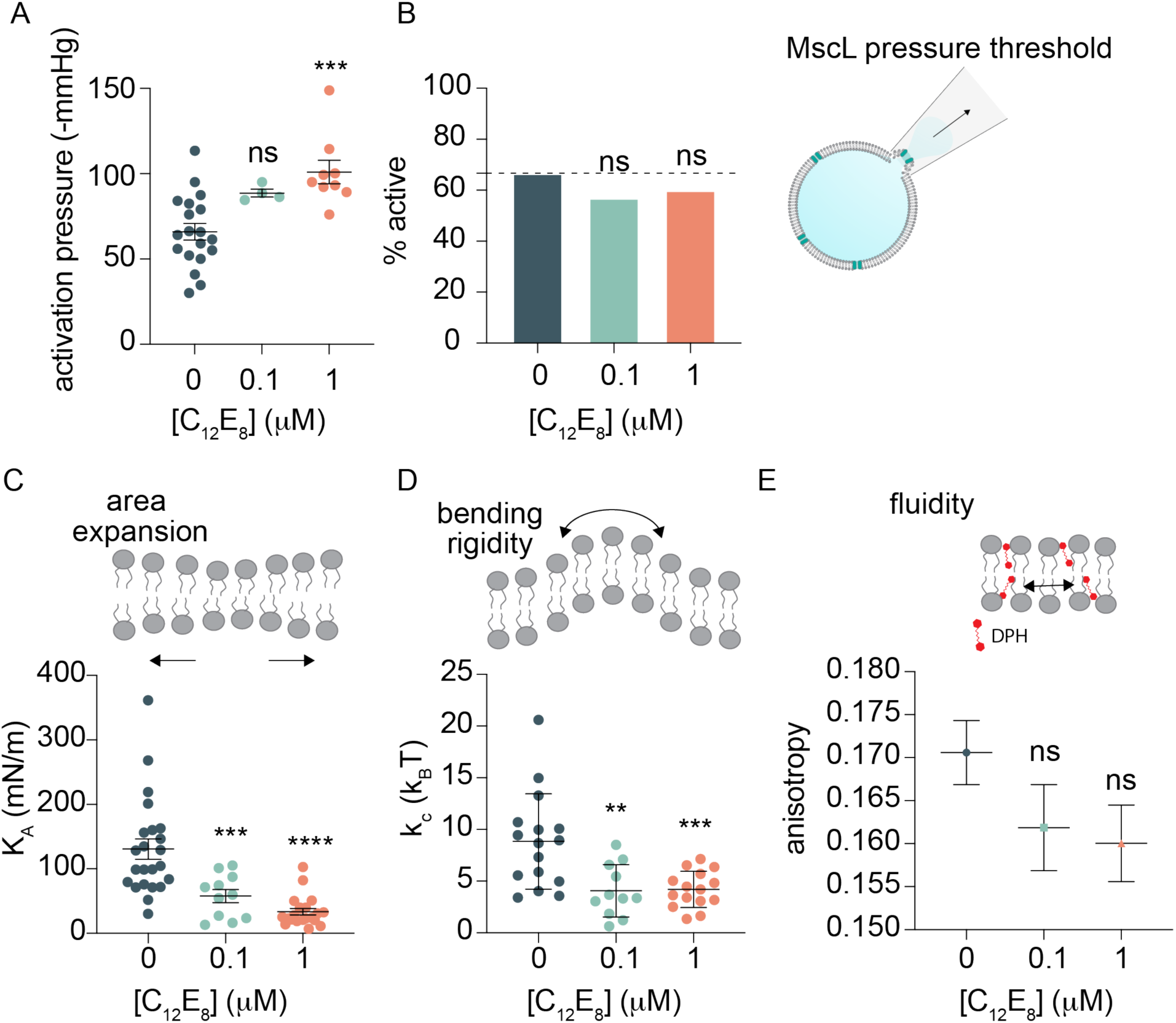
C12E8 effect on MscL activation pressure, K_A_, k_c_, and fluidity. A) C12E8 increases MscLG22S activation pressure with increasing concentration. MscLG22S activation pressure in the presence of C12E8 in the buffer during GPMV formation was measured using patch-clamp electrophysiology and quantified as the pressure at the first channel activation. B) C12E8 did not prevent MscLG22S activation to the same extent as P124 (Figure 2 – figure supplement 1). The percent of recordings which activated MscLG22S were quantified. p-values were generated using Fisher’s exact test compared to 0% C12E8. non-significant (ns) p > 0.05, n > 10. C12E8 decreased K_A_ (C), k_c_ (D) as measured by micropipette aspiration. Points represent individual GPMV measurements which were not repeated. E) C12E8 noticeably but non-significantly increased membrane fluidity as measured by fluorescence anisotropy, where a decreased anisotropy indicates a relative increase in membrane fluidity. Mean and standard error of the mean for all plots is indicated by the bar and error bars p-values in (A), (C-E) were generated by ANOVA using Dunnett’s test of multiple comparisons compared to no C12E8. **** p ≤ 0.0001, *** p ≤ 0.001, ** p ≤ 0.01, non-significant (ns) p >0.05, n > 10.

**Figure 3–figure supplement 2.**
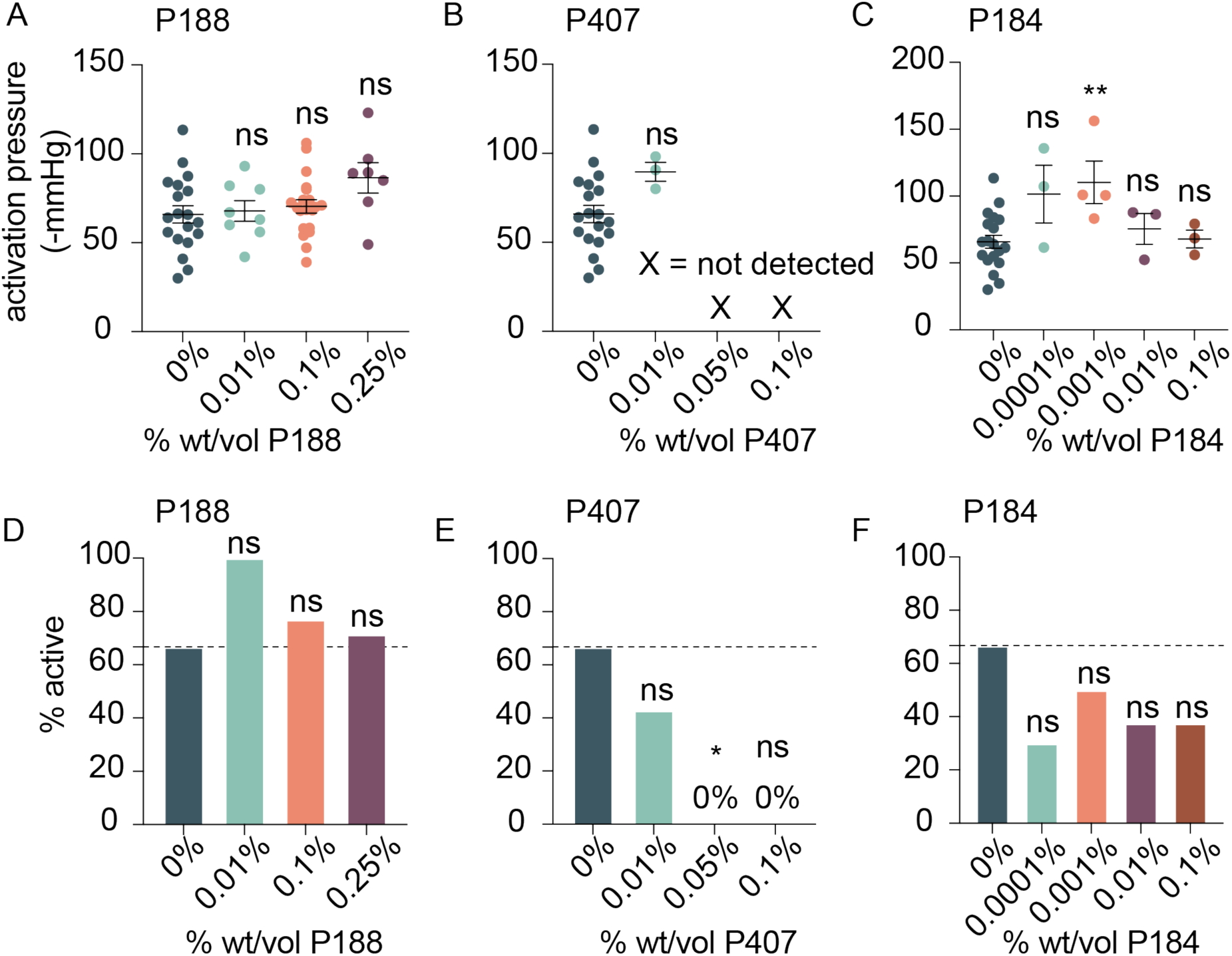
MscL activation sensitivity in the presence of various alternate poloxamers. A-C) MscLG22S activation pressure is altered by poloxamer identity and concentration. The pressure of MscLG22S first channel opening was calculated in the presence of various poloxamers at increasing concentration of poloxamer in the buffer P188 (A), P407 (B), and P184 (C). Individual points represent independent GPMV recordings. Higher concentrations of P407 (B) destabilized the membrane and led to rupture at pressures lower than the expected activation pressure of MscLG22S. For such conditions, no active MscLG22S currents were observed in the small number of P407 GPMVs which were successfully recorded. Mean and standard error of the mean are indicated by the black bar and error bars. The amount of poloxamer in the GPMV formation buffer is indicated in the plot. p-values were generated using ANOVA Dunnett’s test for multiple comparisons compared to no poloxamer (0%). ** p ≤ 0.01, non-significant (ns) p > 0.05, n > 10 for all samples except for P407, the inclusion of which led to membrane instability which prevented MscLG22S activation (B). D-E) MscLG22S activation percentage is modulated by poloxamers. P188 (D) had no depression effect on MscLG22S activation percentage while P407 (E) demonstrated a significant reduction in activation probability. P184 (F) showed a noticeable, but non-significant reduction in MscLG22S activation probability. p-values were generated using Fisher’s exact test compared to 0% poloxamer. * p ≤ 0.05, non-significant (ns) p > 0.05, n > 3.

**Figure 3–figure supplement 3.**
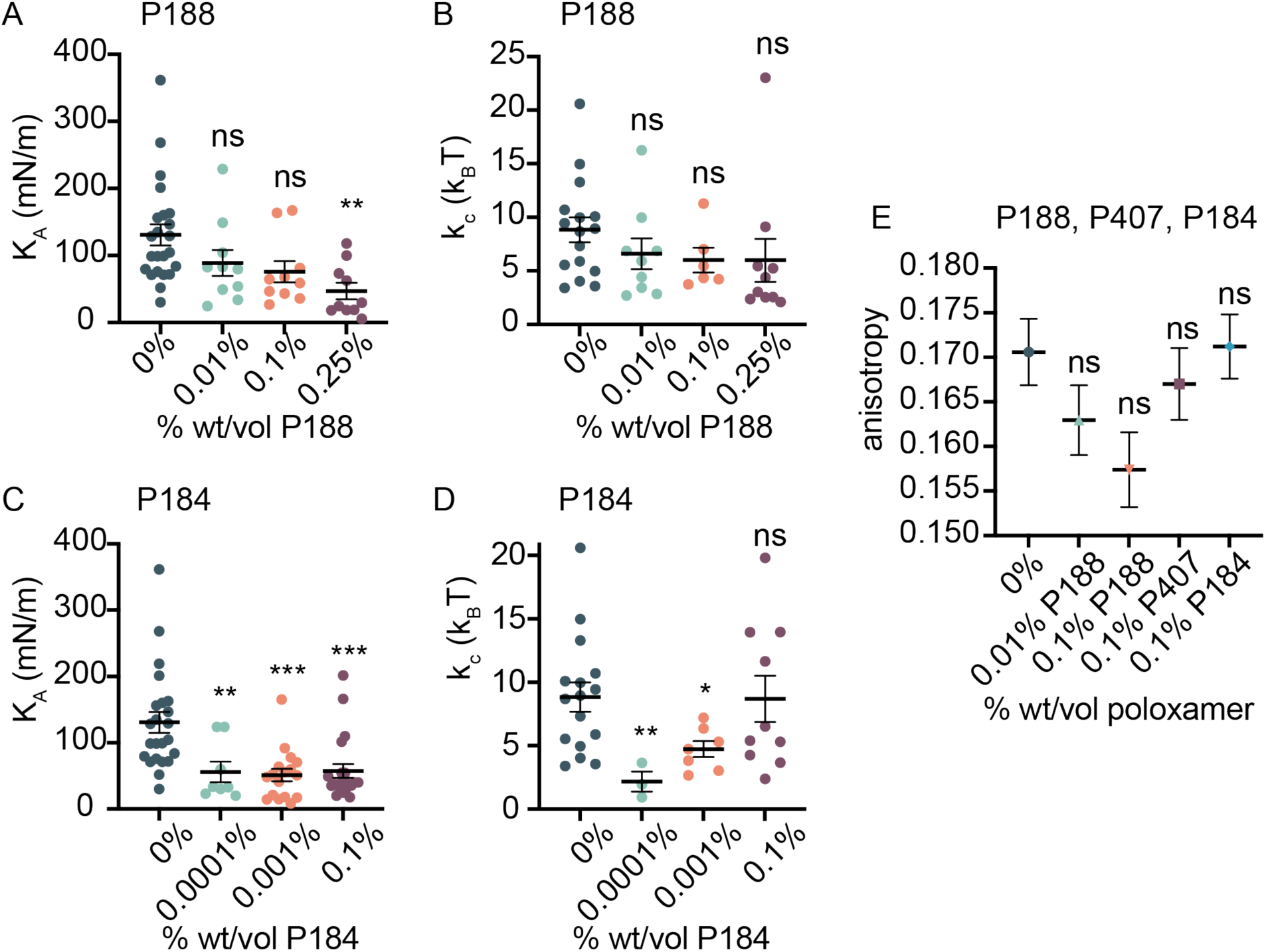
Other poloxamers effect on K_A_, k_c_, and fluidity. Membrane mechanical properties were measured in the presence of poloxamer molecules P188 and P184. A, B) With increasing P188 concentration, K_A_ is reduced and k_c_ is slightly but non-significantly reduced. C) P184 generally decreases membrane K_A_ at and above 0.0001% wt/vol in the GPMV formation buffer. D) P184 exhibits a non-monotonic effect on membrane k_c_ where low concentrations decrease k_c_, but higher concentrations increase k_c_. E) membrane anisotropy was measured in the presence of P188, P407 and P184. No significant differences in membrane fluidity were observed in the presence of these poloxamer molecules. Decreased anisotropy indicates a relative increase in membrane fluidity. Individual points represent single GPMV measurements, mean and standard error of the mean are plotted as a black bar with error bars, pH 7.4. p-values were generated using ANOVA Dunnett’s test for multiple comparisons compared to no poloxamer (0%). *** p ≤ 0.001, ** p ≤ 0.01, * p ≤ 0.05, non-significant (ns) p >0.05, n > 10.

**Figure 3–figure supplement 4.**
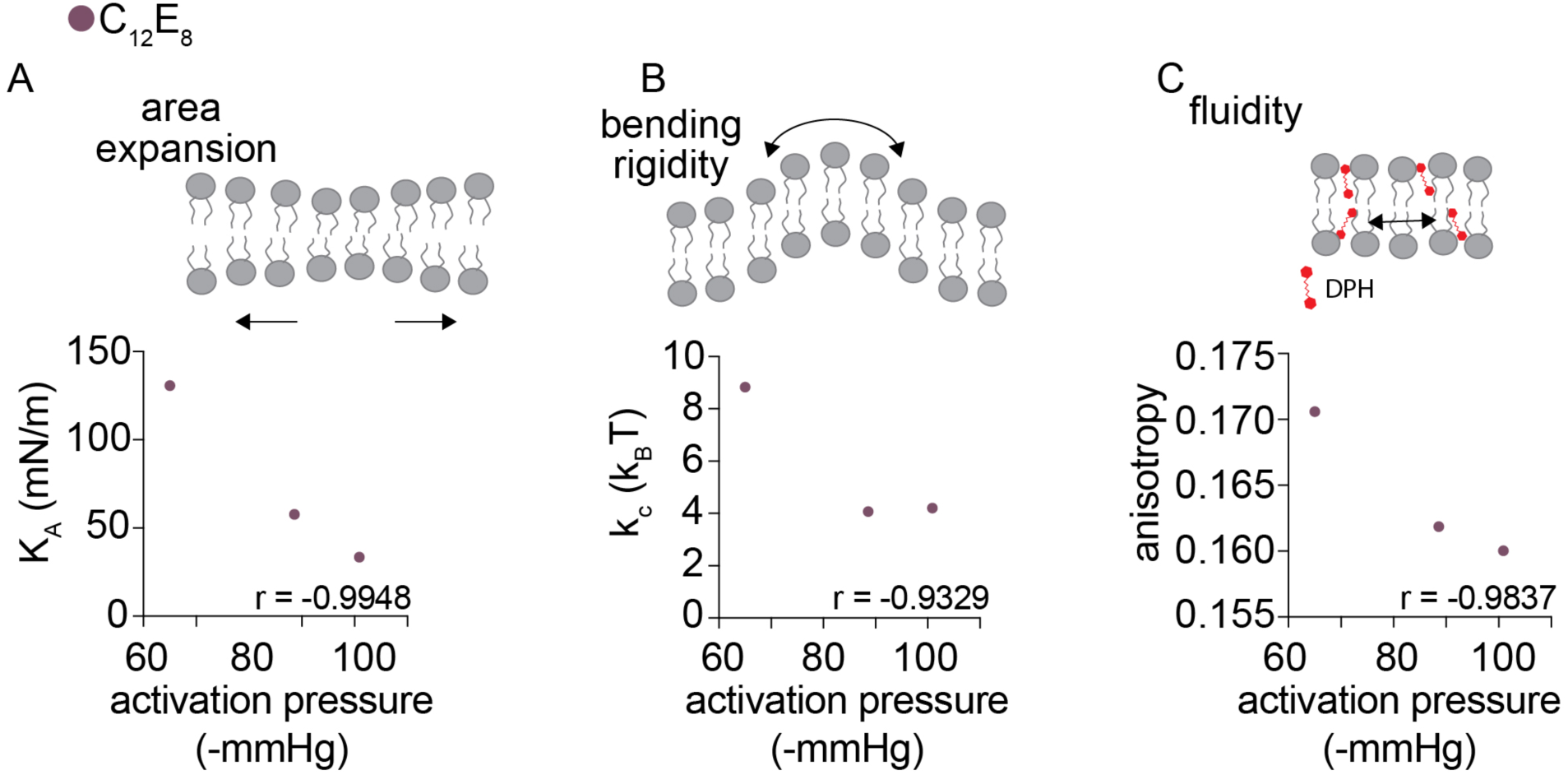
A non-ionic detergent decreases MscL sensitivity and alters membrane properties similarly to poloxamer. A) C12E8 decreases area expansion modulus and increases MscLG22S activation pressure (Figure 3-figure supplement 1A,C). K_A_ was measured using micropipette aspiration. Mean K_A_ is plotted as a function of mean MscLG22S activation pressure for each concentration of C12E8. B) C12E8 decreases membrane k_c_ and increases MscLG22S activation pressure (Figure 3-figure supplement 1A,D). Mean k_c_ is plotted as a function of mean MscLG22S activation pressure. C) Membrane fluidity and MscLG22S activation pressure positively correlate in the presence of C12E8. Mean fluorescence anisotropy of DPH is plotted as a function of MscLG22S activation pressure. Decreased anisotropy indicates a relative increase in membrane fluidity. For all plots, Pearson’s r was calculated to measure correlation strength, n>10.

**Figure 4–figure supplement 1.**
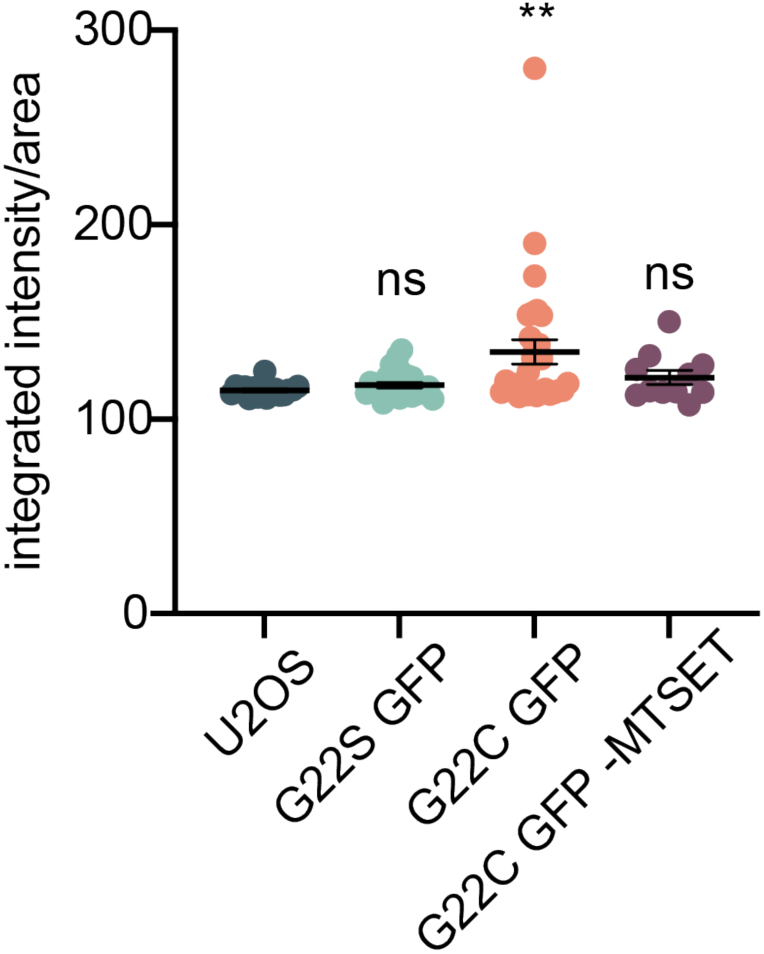
Validation of GPMV calcium influx assay. MscLG22C is a chemically-activatable mutant of MscL which opens in the presence of MTSET through binding to cysteine groups which induces forced hydration of the hydrophobic gate. Calcium and MTSET were added to GPMVs formed from cells incubated with Fluo4 Ca^2+^ indicator and Ca^2+^ influx was detected as an increase in fluorescence and was quantified as integrated intensity/area. GPMVs formed from non-transfected cells (U2OS), MscLG22S-containing GPMVs treated with MTSET, and MscLG22C-containing GPMVs not treated with MTSET (G22CGFP(-MTSET)) did not exhibit increases in fluorescence in response to Ca^2+^. p-values were generated by ANOVA compared to no MscL (U2OS). ** p ≤ 0.01, non-significant (ns), p > 0.05, n > 10 independent GPMVs.

**Figure 4–figure supplement 2.**
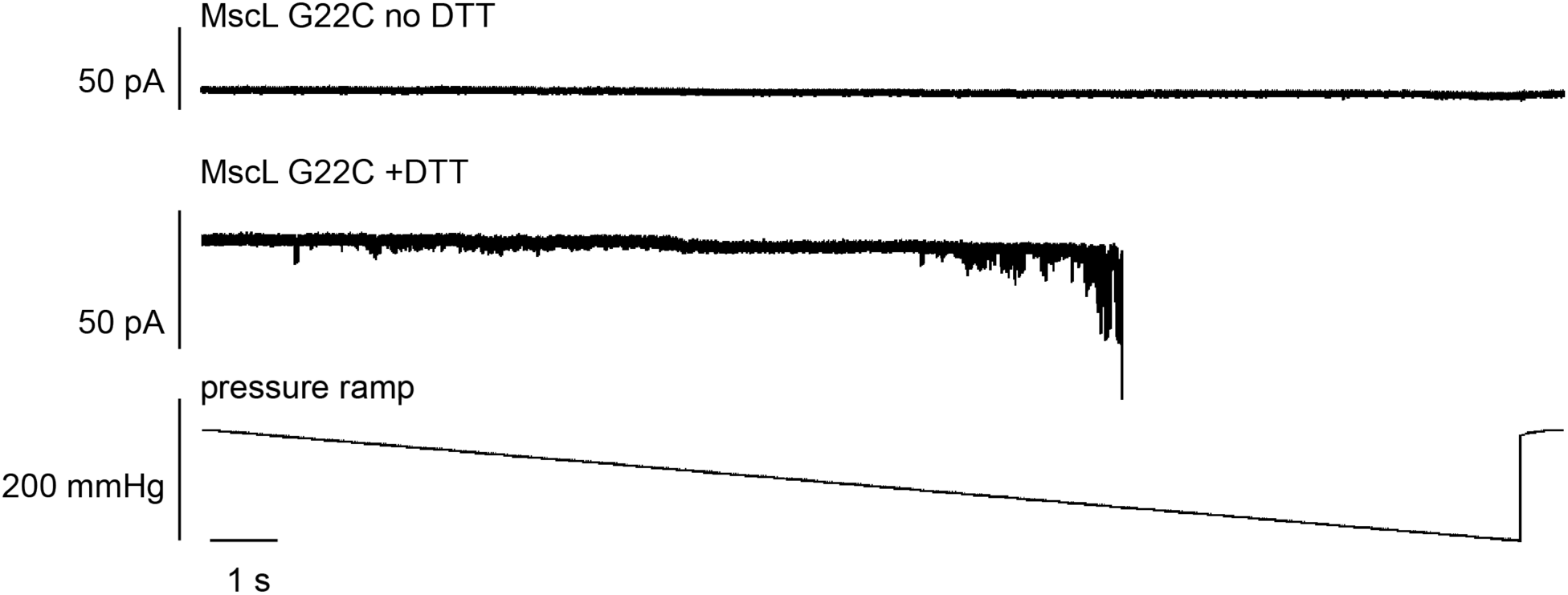
MscLG22C pressure sensitivity is restored in the presence of DTT. The chemically activatable mutant, MscLG22C, used in Ca^2+^ imaging assays was further characterized using patch-clamp electrophysiology to confirm channel activity. Without DTT (top) the channel is not mechanically activatable as described previously (Levin and Blount, 2004) due to disulfide bonds which prevent channel opening. In the presence of DTT, channel mechanosensitivity is restored and stochastic channel activation is observed with increasing pressure.

**Figure 4–figure supplement 3.**
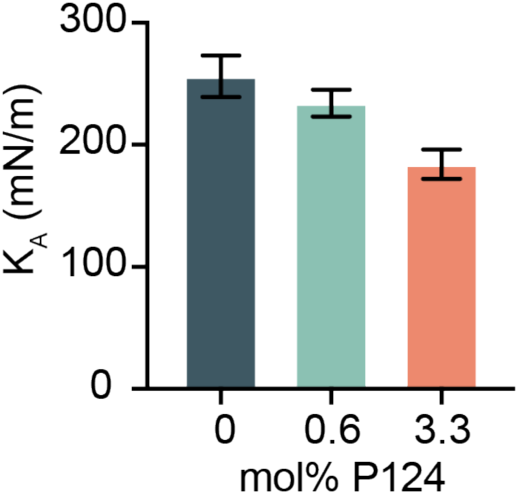
Coarse-grain molecular dynamics simulations confirm the effect of P124 on membrane elasticity. Membrane K_A_ was measured at increasing concentrations of P124 and was found to decrease similar to experimental results. Error bars represent standard error of the mean, n=3.

**Figure 5–figure supplement 1.**
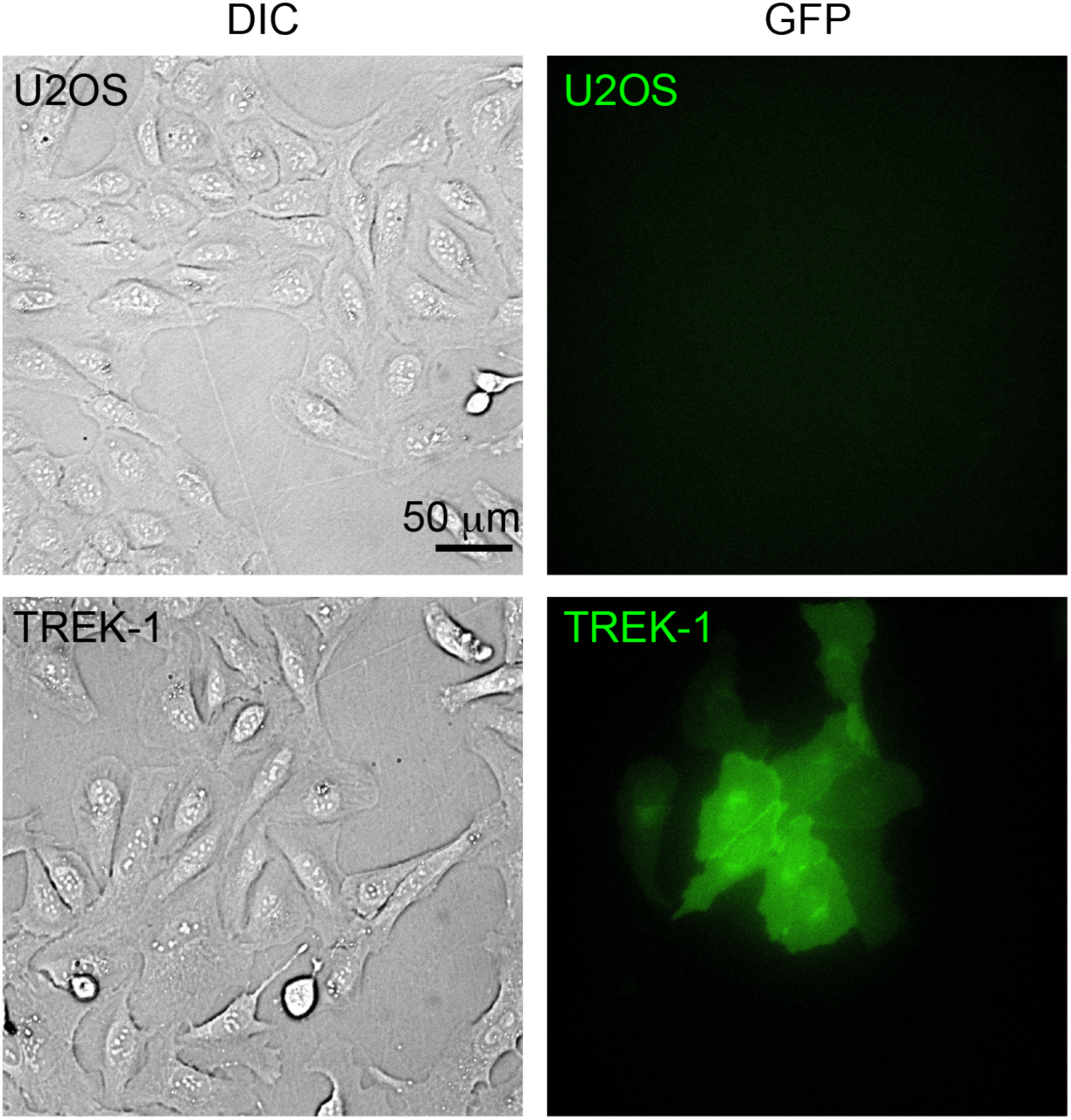
U2OS cells stably expressing mTREK-1. mTREK-1 tagged with GFP localizes to the membrane as shown in GFP images for transfected cells (bottom). Non-transfected U2OS lines (top) do not show GFP signal. DIC images are shown to demonstrate cell boundary and confluency.

**Figure 5–figure supplement 2.**
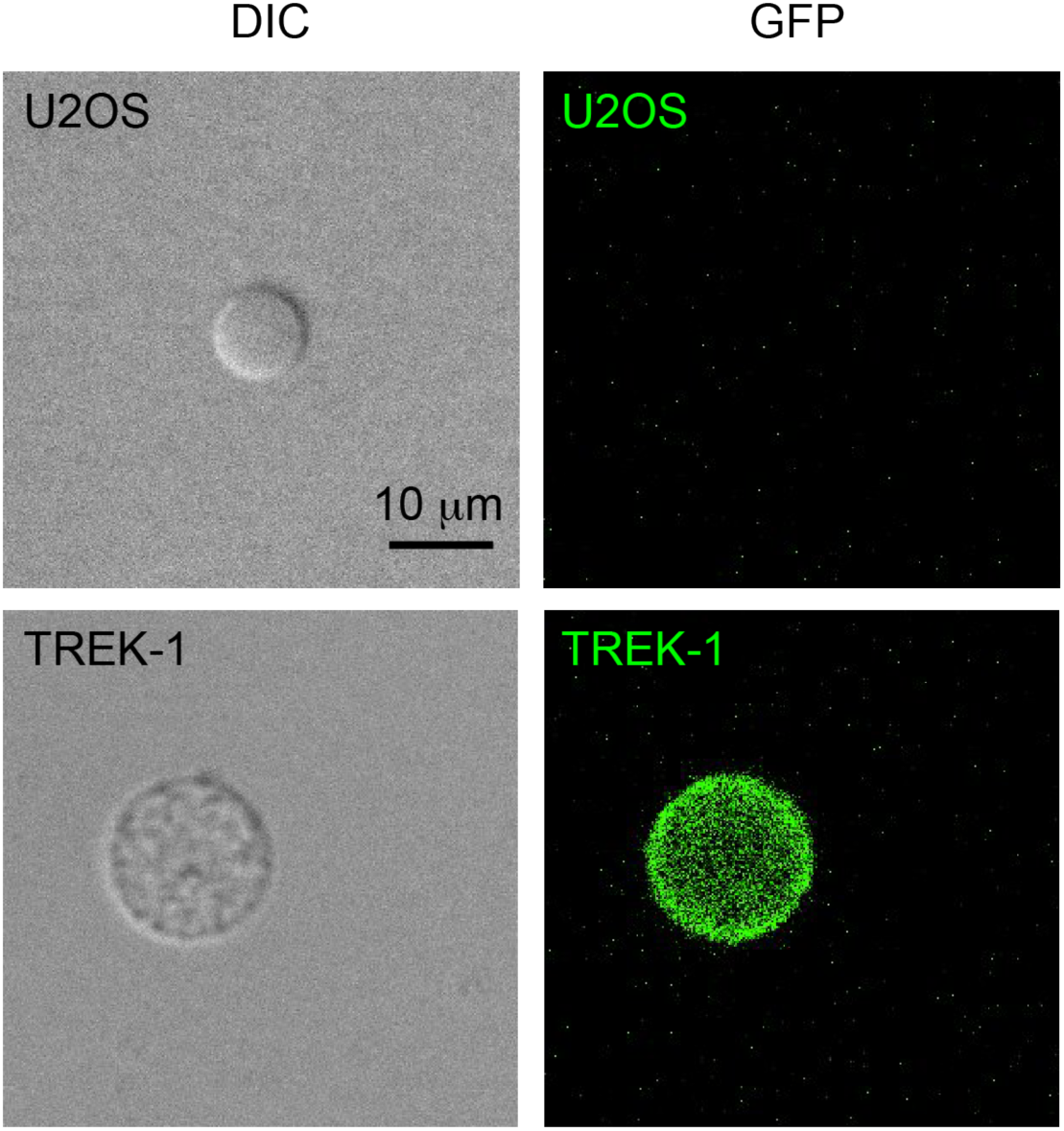
TREK-1 is retained in GPMV membranes. TREK-1 tagged with GFP is localized to the GPMV membrane in transfected cells (bottom). No GFP signal is detected in GPMVs from non-transfected cells (U2OS, top).

**Figure 5–figure supplement 3.**
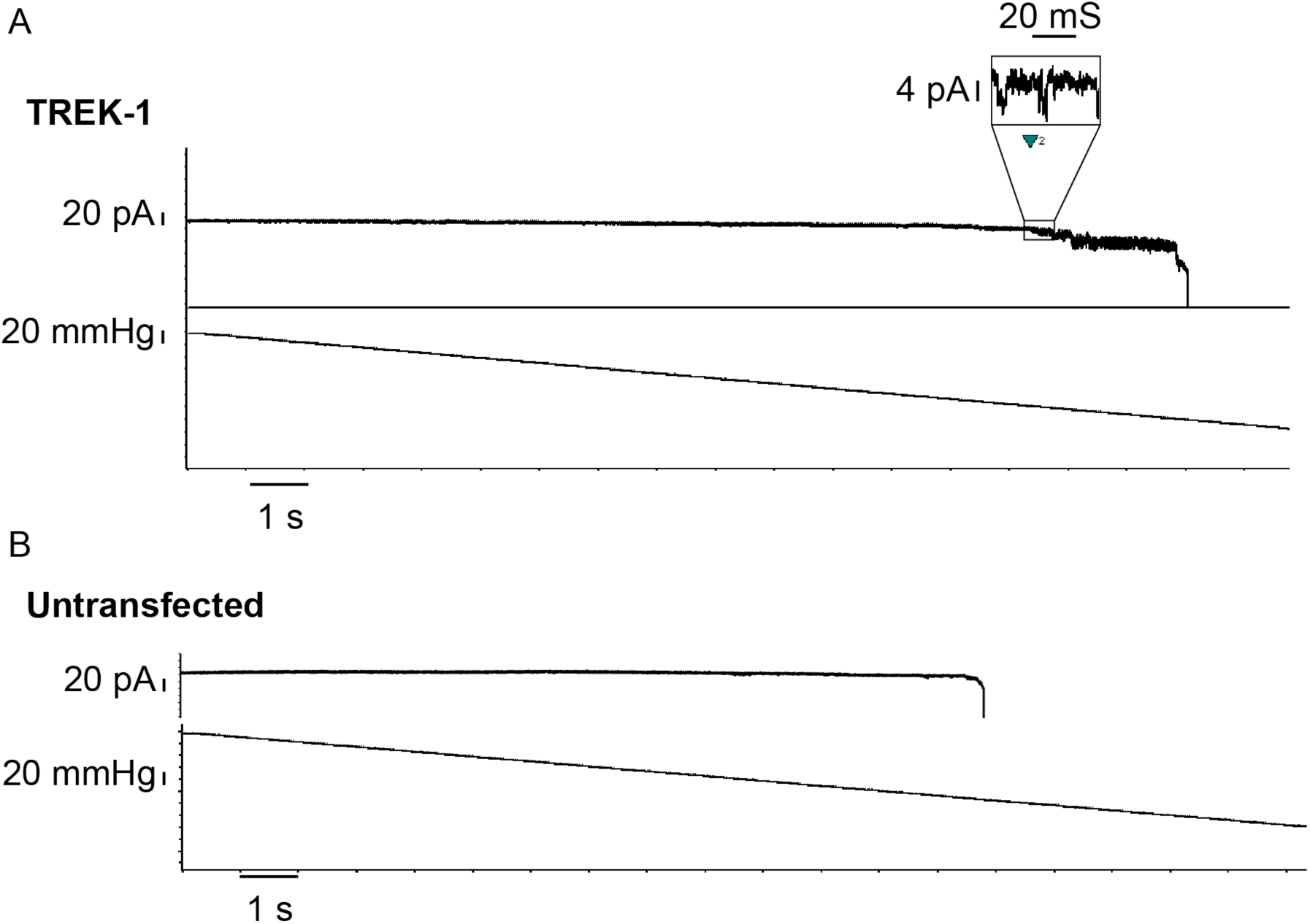
TREK-1 mechanosensitive currents are only observed in GPMVs formed from stable cells expressing TREK-1. A) We observed TREK-1 activation where single channel currents were ∼4.5 pA at -50 mV. TREK-1 activation was observed in most patches above -30 mmHg. B) To further confirm that we observed TREK-1 channels in our electrophysiology recordings, we performed patch-clamp electrophysiology on GPMVs formed from U2OS cells that were untransfected, referred to as untransfected GPMVs, under identical conditions as our TREK-1 recordings which required higher holding potentials to observe single channels due to the smaller conductance of TREK-1 compared to MscLG22S. We applied enough pressure to pop the GPMV membrane to ensure the pressure was high enough to activate any native mechanosensitive channels in the GPMVs. We did not observe any channels in untransfected GPMVs before membrane popping. Recordings are performed at -50 mV in asymmetric potassium buffer conditions. Bath solution was composed of 10 mM HEPES, 150 mM KCl, 2 mM MgCl2 for excised-patch recordings on TREK-1 with outward rectifying potassium currents. n > 6 recordings on independent patches.

## Supplementary Notes

### Supplementary Note 1

The following is a continuation of the thermodynamic model of MscL opening presented in the Discussion. Remember that Δ*G* refers to the free energy cost of opening the channel per the below equation.

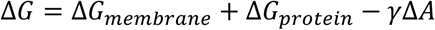

Recall that the term Δ*G*_*membrane*_ refers to the energetic costs incurred from the elastic deformations of the lipid bilayer that are required to accommodate MscL, Δ*G*_*protein*_ refers to the energy difference of the open and closed state of MscL, and −*γ*Δ*A* describes the impetus for channel activation which is the product of the protein surface area gained (Δ*A*) by opening the channel with applied tension (*γ*). The primary focus of the Discussion was the −*γ*Δ*A* term and the Δ*G*_*membrane*_ term; here, we explain key assumptions and further rationale for assuming Δ*G*_*membrane*_ and Δ*G*_*protein*_ are not substantial contributors when explaining the different behaviors of MscL opening in membranes with and without poloxamer 124.

Beginning with important assumptions, we note that we experimentally investigated how MscL opening pressure varies with membrane composition (e.g., Figures 2B, 3) using a patch-clamp setup with an integrated pressure transducer. At constant micropipette and GPMV geometry in such an experiment, one can assume that the applied channel-opening pressure (*P*_*open*_) is proportional to the opening free energy of MscL (Δ*G*) (Wiggins and Phillips, 2005). We will further assume that changes in opening pressure for MscL in different membrane compositions (Δ*P*_*open*_) is proportional to changes in free energy of MscL opening in those compositions (ΔΔ*G*_*exp*_, i.e., the free energy change from a lipid-only membrane to a polymer-containing membrane).

In line with other work, we assume that Δ*G*_*protein*_ remains constant in experiments where we modified membrane compositions using the same MscL variants to observe ΔΔ*G*_*exp*_ (Melo et al., 2017; Wiggins and Phillips, 2005). Indeed, it has been shown that MscL opening via delipidation of the transmembrane pocket and via applied tension results in the same channel structures for very different pore-opening forces (B. Wang et al., 2021). This supports our use of a constant Δ*G*_*protein*_ in our experiments for tension-mediated pore opening for a given MscL variant. We also assume the lipid-protein interaction energy between the open and closed states for membranes with and without P124 are similar. This assumption is consistent with other reports using alcohols to modify membranes (Melo et al., 2017) and is supported by the observations in Figure 4A-E that P124 does not impact conductance and therefore likely does not occlude the channel pore. Therefore, although we cannot rule out the possibly that protein-membrane interactions may change slightly with the incorporation of P124 into membranes, our assumption that Δ*G*_*protein*_ remains constant is supported by our observations and other reports.

The contribution Δ*G*_*membrane*_can be further dissected into several terms that, in the typically used continuum mechanical membrane model, depend on the membrane elastic parameters of bending rigidity, area expansion modulus and membrane spontaneous curvature (Phillips et al., 2009). To date, hydrophobic thickness and local curvature strain have emerged as the predominant such modulators of MscL activation. In general, thinner membranes reduce the activation pressure of MscL as a result of hydrophobic mismatch (Katsuta et al., 2019; Ridone et al., 2018) and high local curvature favors the channel open state (Bavi et al., 2014; Doerner et al., 2012; Nomura et al., 2012; Wang et al., 2014). In the following two paragraphs, we are considering differences in membrane properties resulting from the addition of polymer to membranes.

At a high level, we expect Δ*G*_*membrane*_ to penalize the open MscL configuration in part because a typical membrane hydrophobic thickness is larger than the hydrophobic region of the open, “thinned”, MscL configuration—this difference can induce a large hydrophobic mismatch between membrane thickness and channel thickness that is thermodynamically unfavorable (Perozo et al., 2002; Phillips et al., 2009; Ursell et al., 2007; Wiggins and Phillips, 2005). From simulations (Figure 4I), we found only very small changes in membrane thickness with polymer P124, suggesting that the Δ*G*_*membrane*_ cost due to hydrophobic mismatch is highly similar for membranes with and without P124.

With respect to Δ*G*_*membrane*_, it is important to consider membrane spontaneous curvature which results from membrane asymmetry (Perozo et al., 2002). This effect is highlighted in experiments where LPC was added to the GPMVs from the outside—here, we expect asymmetry to initially exist. As a result, the Δ*G*_*membrane*_ energetic cost of channel opening is lowered because asymmetric LPC incorporation drives membrane curvature changes that favor opening of the MscL pore (Perozo et al., 2002; Yoo and Cui, 2009). Over longer timescales, we expect LPC to be equilibrated between leaflets, and the LPC-mediated reduction of activation sensitivity vanishes (Figure 2-figure supplement 7B). Similarly, we exclude the contribution from spontaneous curvature when considering experiments with C12E8 and polymer compared to lipid-only membranes because the additive’s distribution is likely equilibrated between bilayer leaflets during the patch clamp experiments. Equilibration is facilitated by the hour-long incubation times and perhaps by enhanced permeability of GPMVs (Skinkle, Levental, and Levental 2020). Such equilibration is also consistent with fast equilibration times of poloxamer observed using supported lipid bilayers (Kim et al. 2020). We conclude that membrane asymmetry is not a substantial factor for Δ*G*_*membrane*_ differences between membranes with and without polymer.

In the Discussion, we reason that the magnitude of ΔΔ*G*_*membrane*_ arising from changes in K_A_ and K_c_ is smaller than the differences in *γ*Δ*A* between membrane compositions (∼50-79 KbT). Here we provide order of magnitude estimates for ΔΔ*G*_*membrane*_ from literature and our results. The cost of a midplane bilayer deformation due to channel opening is small (< 1 KbT) and scales with K_c_^1/2^, and the cost to deform the bilayer thickness is moderate (∼10 KbT) and scales with K_c_^1/4^ * K_A_^3/4^ (Ollila et al., 2011; Wiggins and Phillips, 2005). The former term is small enough to omit from this analysis. From Figure 2E,F, K_A_ and K_c_ decreased by ∼3.8x and ∼2.3x, respectively, in the presence of high concentrations of P124; we approximate the ΔΔ*G*_*membrane*_ term is reduced ∼3.4x to ∼3 KbT, much smaller than our 50-79 KbT estimate for ΔΔ*G*_*exp*_.

## Notes

### Competing Interest Statement

The authors have declared no competing interest.

### Summary of Updates

We have performed additional simulations and expanded our discussion to compare our data with a broader range of existing models. Briefly, we show that interfacial tension, a membrane property directly related to membrane area expansion and rigidity, is a significant driver for MscL channel activation. Accordingly, our results support the force-from-lipid hypothesis in a new way: by directly toggling this parameter (interfacial tension), we can drive channel activation.

## References

Anishkin A, Loukin SH, Teng J, Kung C. 2014. Feeling the hidden mechanical forces in lipid bilayer is an original sense. Proc Natl Acad Sci 111:7898–7905. doi:10.1073/pnas.1313364111

Balleza D. 2012. Mechanical properties of lipid bilayers and regulation of mechanosensitive function: From biological to biomimetic channels. Channels 6:220–233. doi:10.4161/chan.21085

Banerjee A, Lee A, Campbell E, MacKinnon R. 2013. Structure of a pore-blocking toxin in complex with a eukaryotic voltage-dependent K+ channel. Elife 2013:1–22. doi:10.7554/eLife.00594

Battaglia G, Ryan AJ. 2005. Bilayers and interdigitation in block copolymer vesicles. J Am Chem Soc 127:8757–8764. doi:10.1021/ja050742y

Bavi N, Cortes DM, Cox CD, Rohde PR, Liu W, Deitmer JW, Bavi O, Strop P, Hill AP, Rees D, Corry B, Perozo E, Martinac B. 2016. The role of MscL amphipathic N terminus indicates a blueprint for bilayer-mediated gating of mechanosensitive channels. Nat Commun 7. doi:10.1038/ncomms11984

Bavi O, Vossoughi M, Naghdabadi R, Jamali Y. 2014. The effect of local bending on gating of MscL using a representative volume element and finite element simulation. Channels 8:344–349. doi:10.4161/chan.29572

Bermudez H, Brannan AK, Hammer DA, Bates FS, Discher DE. 2002. Molecular weight dependence of polymersome membrane structure, elasticity, and stability. Macromolecules 35:8203–8208. doi:10.1021/ma020669l

Brohawn SG, Su Z, MacKinnon R. 2014. Mechanosensitivity is mediated directly by the lipid membrane in TRAAK and TREK1 K ^+^ channels. Proc Natl Acad Sci 111:3614–3619. doi:10.1073/pnas.1320768111

Brohawn SG, Wang W, Handler A, Campbell EB, Schwarz JR, MacKinnon R. 2019. The mechanosensitive ion channel TRAAK is localized to the mammalian node of Ranvier. Elife 8:1–22. doi:10.7554/eLife.50403

Bruno MJ, Koeppe RE, Andersen OS. 2007. Docosahexaenoic acid alters bilayer elastic properties. Proc Natl Acad Sci 104:9638–9643. doi:10.1073/pnas.0701015104

Cahalan SM, Lukacs V, Ranade SS, Chien S, Bandell M, Patapoutian A. 2015. Piezo1 links mechanical forces to red blood cell volume. Elife 4. doi:10.7554/eLife.07370

Caires R, Sierra-Valdez FJ, Millet JRM, Herwig JD, Roan E, Vásquez V, Cordero-Morales JF. 2017. Omega-3 Fatty Acids Modulate TRPV4 Function through Plasma Membrane Remodeling. Cell Rep 21:246–258. doi:10.1016/j.celrep.2017.09.029

Calori IR, Pinheiro L, Braga G, Morais FAP de, Caetano W, Tedesco AC, Hioka N. 2022. Interaction of triblock copolymers (Pluronics®) with DMPC vesicles: a photophysical and computational study. Spectrochim Acta Part A Mol Biomol Spectrosc 121178. doi:10.1016/j.saa.2022.121178

Chang G, Spencer RH, Lee AT, Barclay MT, Rees DC. 1998. Structure of the MscL homolog from Mycobacterium tuberculosis: A gated mechanosensitive ion channel. Science (80-) 282:2220–2226. doi:10.1126/science.282.5397.2220

Chiang CS, Anishkin A, Sukharev S. 2004. Gating of the Large Mechanosensitive Channel in Situ: Estimation of the Spatial Scale of the Transition from Channel Population Responses. Biophys J 86. doi:10.1016/S0006-3495(04)74337-4

Choi YK, Park SJ, Park S, Kim S, Kern NR, Lee J, Im W. 2021. CHARMM-GUI Polymer Builder for Modeling and Simulation of Synthetic Polymers. J Chem Theory Comput 17:2431– 2443. doi:10.1021/acs.jctc.1c00169

Cohen AE, Shi Z. 2020. Do Cell Membranes Flow Like Honey or Jiggle Like Jello? BioEssays 42:1–13. doi:10.1002/bies.201900142

Cordero-Morales JF, Vásquez V. 2018. How lipids contribute to ion channel function, a fat perspective on direct and indirect interactions. Curr Opin Struct Biol 51:92–98. doi:10.1016/j.sbi.2018.03.015

Cox CD, Bavi N, Martinac B. 2019. Biophysical Principles of Ion-Channel-Mediated Mechanosensory Transduction. Cell Rep 29:1–12. doi:10.1016/j.celrep.2019.08.075

Dave N, Cetiner U, Arroyo D, Fonbuena J, Tiwari M, Barrera P, Lander N, Anishkin A, Sukharev S, Jimenez V. 2021. A novel mechanosensitive channel controls osmoregulation, differentiation, and infectivity in trypanosoma cruzi. Elife 10:1–32. doi:10.7554/ELIFE.67449

del Mármol JI, Touhara KK, Croft G, MacKinnon R. 2018. Piezo1 forms a slowly-inactivating mechanosensory channel in mouse embryonic stem cells. Elife 7:1–18. doi:10.7554/eLife.33149

Doerner JF, Febvay S, Clapham DE. 2012. Controlled delivery of bioactive molecules into live cells using the bacterial mechanosensitive channel MscL. Nat Commun 3:990. doi:10.1038/ncomms1999

Evans EA, Waugh R, Melnik L. 1976. Elastic area compressibility modulus of red cell membrane. Biophys J 16:585–595. doi:10.1016/S0006-3495(76)85713-X

Feller SE, Pastor RW. 1999. Constant surface tension simulations of lipid bilayers: The sensitivity of surface areas and compressibilities. J Chem Phys 111:1281–1287. doi:10.1063/1.479313

Fleury JB, Baulin VA. 2021. Microplastics destabilize lipid membranes by mechanical stretching. Proc Natl Acad Sci U S A 118:1–8. doi:10.1073/pnas.2104610118

Flyvbjerg H, Petersen HG. 1989. Error estimates on averages of correlated data. J Chem Phys 91:461–466. doi:10.1063/1.457480

Goodman KE, Hare JT, Khamis ZI, Hua T, Sang QXA. 2021. Exposure of Human Lung Cells to Polystyrene Microplastics Significantly Retards Cell Proliferation and Triggers Morphological Changes. Chem Res Toxicol 34:1069–1081. doi:10.1021/acs.chemrestox.0c00486

Grage SL, Afonin S, Ieronimo M, Berditsch M, Wadhwani P, Ulrich AS. 2022. Probing and Manipulating the Lateral Pressure Profile in Lipid Bilayers Using Membrane-Active Peptides—A Solid-State 19F NMR Study. Int J Mol Sci 23:4544. doi:10.3390/ijms23094544

Großkopf S, Fouquet P, Wiehemeier L, Hellweg T. 2021. Pluronic-based lamellar phases: influence of polymer architecture on bilayer bending elasticity. Mol Phys 119. doi:10.1080/00268976.2021.1893400

Guo YR, MacKinnon R. 2017. Structure-based membrane dome mechanism for piezo mechanosensitivity. Elife 6:1–19. doi:10.7554/eLife.33660

Guzniczak E, Jimenez M, Irwin M, Otto O, Willoughby N, Bridle H. 2018. Impact of poloxamer 188 (Pluronic F-68) additive on cell mechanical properties, quantification by real-time deformability cytometry. Biomicrofluidics 12:044118. doi:10.1063/1.5040316

Heureaux J, Chen D, Murray VL, Deng CX, Liu AP. 2014. Activation of a Bacterial Mechanosensitive Channel in Mammalian Cells by Cytoskeletal Stress. Cell Mol Bioeng 7:307–319. doi:10.1007/s12195-014-0337-8

Heurteaux C, Guy N, Laigle C, Blondeau N, Duprat F, Mazzuca M, Lang-Lazdunski L, Widmann C, Zanzouri M, Romey G, Lazdunski M. 2004. TREK-1, a K+ channel involved in neuroprotection and general anesthesia. EMBO J 23:2684–2695. doi:10.1038/sj.emboj.7600234

Hollóczki O, Gehrke S. 2019. Can Nanoplastics Alter Cell Membranes? ChemPhysChem 9–12. doi:10.1002/cphc.201900481

Houk AR, Jilkine A, Mejean CO, Boltyanskiy R, Dufresne ER, Angenent SB, Altschuler SJ, Wu LF, Weiner OD. 2012. Membrane tension maintains cell polarity by confining signals to the leading edge during neutrophil migration. Cell 148:175–188. doi:10.1016/j.cell.2011.10.050

Huang SK, Almurad O, Pejana RJ, Morrison ZA, Pandey A, Picard LP, Nitz M, Sljoka A, Prosser RS. 2022. Allosteric modulation of the adenosine A2A receptor by cholesterol. Elife 11:1– 24. doi:10.7554/ELIFE.73901

Jacobs ML, Boyd MA, Kamat NP. 2019. Diblock copolymers enhance folding of a mechanosensitive membrane protein during cell-free expression. Proc Natl Acad Sci 116:4031–4036. doi:10.1073/pnas.1814775116

Jacobs ML, Faizi HA, Peruzzi JA, Vlahovska PM, Kamat NP. 2021. EPA and DHA differentially modulate membrane elasticity in the presence of cholesterol. Biophys J 120:2317–2329. doi:10.1016/j.bpj.2021.04.009

Jo S, Kim T, Iyer GV, Im W. 2008. CHARMM-GUI: A Web-Based Graphical User Interface for CHARMM. J Comput Chem 29:1859–1865. doi:10.1002/jcc

Johnson SL. 2015. Membrane properties specialize mammalian inner hair cells for frequency or intensity encoding. Elife 4:1–21. doi:10.7554/elife.08177

Katsuta H, Sawada Y, Sokabe M. 2019. Biophysical Mechanisms of Membrane-Thickness-Dependent MscL Gating: An All-Atom Molecular Dynamics Study. Langmuir 35:7432– 7442. doi:10.1021/acs.langmuir.8b02074

Keller H, Lorizate M, Schwille P. 2009. PI(4,5)P2 Degradation Promotes the Formation of Cytoskeleton-Free Model Membrane Systems. ChemPhysChem 10:2805–2812. doi:10.1002/cphc.200900598

Kwok R, Evans E. 1981. Thermoelasticity of large lecithin bilayer vesicles. Biophys J 35:637– 652. doi:10.1016/S0006-3495(81)84817-5

LaMotte RH. 2016. Allergic Contact Dermatitis: A Model of Inflammatory Itch and Pain in Human and Mouse In: Ma C, Huang Y, editors. Translational Research in Pain and Itch. Dordrecht: Springer Netherlands. pp. 23–32. doi:10.1007/978-94-017-7537-3_2

Levental KR, Lorent JH, Lin X, Skinkle AD, Surma MA, Stockenbojer EA, Gorfe AA, Levental I. 2016. Polyunsaturated lipids regulate membrane domain stability by tuning membrane order. Biophys J 110:1800–1810. doi:10.1016/j.bpj.2016.03.012

Levin G, Blount P. 2004. Cysteine Scanning of MscL Transmembrane Domains Reveals Residues Critical for Mechanosensitive Channel Gating. Biophys J 86:2862–2870. doi:10.1016/S0006-3495(04)74338-6

Lewis AH, Grandl J. 2015. Mechanical sensitivity of Piezo1 ion channels can be tuned by cellular membrane tension. Elife 4. doi:10.7554/eLife.12088

Lipowsky R, Sackmann E. 1995. Structure and dynamics of membranes, Handbook of biological physics; v. 1A-B. Amsterdam; Elsevier Science.

Lolicato M, Arrigoni C, Mori T, Sekioka Y, Bryant C, Clark KA, Minor DL. 2017. K2P2.1 (TREK-1)–activator complexes reveal a cryptic selectivity filter binding site. Nature 547:364–368. doi:10.1038/nature22988

Lüchtefeld I, Pivkin I V., Gardini L, Zare-Eelanjegh E, Gäbelein C, Ihle SJ, Reichmuth AM, Capitanio M, Martinac B, Zambelli T, Vassalli M. 2024. Dissecting cell membrane tension dynamics and its effect on Piezo1-mediated cellular mechanosensitivity using force-controlled nanopipettes. Nat Methods 2024 216 21:1063–1073. doi:10.1038/s41592-024-02277-8

Lundbæk JA, Birn P, Hansen AJ, Søgaard R, Nielsen C, Girshman J, Bruno MJ, Tape SE, Egebjerg J, Greathouse D V, Mattice GL, Koeppe RE, Andersen OS. 2004. Regulation of Sodium Channel Function by Bilayer Elasticity: The Importance of Hydrophobic Coupling. Effects of Micelle-forming Amphiphiles and Cholesterol. J Gen Physiol 123:599–621. doi:10.1085/jgp.200308996

Lundbæk JA, Collingwood SA, Ingólfsson HI, Kapoor R, Andersen OS. 2010. Lipid bilayer regulation of membrane protein function: gramicidin channels as molecular force probes. J R Soc Interface 7:373–395. doi:10.1098/rsif.2009.0443

Martinac B, Nikolaev YA, Silvani G, Bavi N, Romanov V, Nakayama Y, Martinac AD, Rohde P, Bavi O, Cox CD. 2020. Cell membrane mechanics and mechanosensory transduction, 1st ed, Current Topics in Membranes. Elsevier Inc. doi:10.1016/bs.ctm.2020.08.002

Melo MN, Arnarez C, Sikkema H, Kumar N, Walko M, Berendsen HJC, Kocer A, Marrink SJ, Ingólfsson HI. 2017. High-Throughput Simulations Reveal Membrane-Mediated Effects of Alcohols on MscL Gating. J Am Chem Soc 139:2664–2671. doi:10.1021/jacs.6b11091

Michaud-Agrawal N, Denning EJ, Woolf TB, Beckstein O. 2011. MDAnalysis: A toolkit for the analysis of molecular dynamics simulations. J Comput Chem 32:2319–2327. doi:10.1002/jcc.21787

Nakayama Y, Komazawa K, Bavi N, Hashimoto K ichi, Kawasaki H, Martinac B. 2018. Evolutionary specialization of MscCG, an MscS-like mechanosensitive channel, in amino acid transport in Corynebacterium glutamicum. Sci Rep 8:1–13. doi:10.1038/s41598-018-31219-6

Nomura T, Cranfield CG, Deplazes E, Owen DM, Macmillan A, Battle AR, Constantine M, Sokabe M, Martinac B. 2012. Differential effects of lipids and lyso-lipids on the mechanosensitivity of the mechanosensitive channels MscL and MscS. Proc Natl Acad Sci 109:8770–8775. doi:10.1073/pnas.1200051109

Ohashi Y, Tremel S, Masson GR, McGinney L, Boulanger J, Rostislavleva K, Johnson CM, Niewczas I, Clark J, Williams RL. 2020. Membrane characteristics tune activities of endosomal and autophagic human VPS34 complexes. Elife 9:1–28. doi:10.7554/eLife.58281

Ollila OHS, Louhivuori M, Marrink SJ, Vattulainen I. 2011. Protein shape change has a major effect on the gating energy of a mechanosensitive channel. Biophys J 100. doi:10.1016/j.bpj.2011.02.027

Otten D, Brown MF, Beyer K. 2000. Softening of Membrane Bilayers by Detergents Elucidated by Deuterium NMR Spectroscopy. J Phys Chem B 104:12119–12129. doi:10.1021/jp001505e

Perozo E, Kloda A, Cortes DM, Martinac B. 2002. Physical principles underlying the transduction of bilayer deformation forces during mechanosensitive channel gating. Nat Struct Biol 9. doi:10.1038/nsb827

Phillips R, Ursell T, Wiggins P, Sens P. 2009. Emerging roles for lipids in shaping membrane-protein function. Nature. doi:10.1038/nature08147

Pliotas C, Naismith JH. 2017. Spectator no more, the role of the membrane in regulating ion channel function. Curr Opin Struct Biol 45:59–66. doi:10.1016/j.sbi.2016.10.017

Procko C, Murthy S, Keenan WT, Mousavi SAR, Dabi T, Coombs A, Procko E, Baird L, Patapoutian A, Chory J. 2021. Stretch-activated ion channels identified in the touch-sensitive structures of carnivorous droseraceae plants. Elife 10:1–24. doi:10.7554/eLife.64250

Rabbel H, Werner M, Sommer JU. 2015. Interactions of Amphiphilic Triblock Copolymers with Lipid Membranes: Modes of Interaction and Effect on Permeability Examined by Generic Monte Carlo Simulations. Macromolecules 48:4724–4732. doi:10.1021/acs.macromol.5b00720

Rawicz W, Olbrich KC, McIntosh T, Needham D, Evans E. 2000. Effect of Chain Length and Unsaturation on Elasticity of Lipid Bilayers. Biophys J 79:328–339. doi:10.1016/S0006-3495(00)76295-3

Reddy B, Bavi N, Lu A, Park Y, Perozo E. 2019. Molecular basis of force-from-lipids gating in the mechanosensitive channel mscs. Elife 8:1–24. doi:10.7554/eLife.50486

Ridone P, Grage SL, Patkunarajah A, Battle AR, Ulrich AS, Martinac B. 2018. “Force-from-lipids” gating of mechanosensitive channels modulated by PUFAs. J Mech Behav Biomed Mater 79:158–167. doi:10.1016/j.jmbbm.2017.12.026

Ridone P, Pandzic E, Vassalli M, Cox CD, Macmillan A, Gottlieb PA, Martinac B. 2020. Disruption of membrane cholesterol organization impairs the activity of PIEZO1 channel clusters. J Gen Physiol 152. doi:10.1085/jgp.201912515

Rokitskaya TI, Nazarov PA, Golovin A V., Antonenko YN. 2017. Blocking of Single α-Hemolysin Pore by Rhodamine Derivatives. Biophys J 112:2327–2335. doi:10.1016/j.bpj.2017.04.041

Romero LO, Massey AE, Mata-Daboin AD, Sierra-Valdez FJ, Chauhan SC, Cordero-Morales JF, Vásquez V. 2019. Dietary fatty acids fine-tune Piezo1 mechanical response. Nat Commun 10:1200. doi:10.1038/s41467-019-09055-7

Schindelin J, Arganda-Carreras I, Frise E, Kaynig V, Longair M, Pietzsch T, Preibisch S, Rueden C, Saalfeld S, Schmid B, Tinevez JY, White DJ, Hartenstein V, Eliceiri K, Tomancak P, Cardona A. 2012. Fiji: An open-source platform for biological-image analysis. Nat Methods. doi:10.1038/nmeth.2019

Sezgin E, Kaiser HJ, Baumgart T, Schwille P, Simons K, Levental I. 2012. Elucidating membrane structure and protein behavior using giant plasma membrane vesicles. Nat Protoc 7:1042–1051. doi:10.1038/nprot.2012.059

Shi Z, Innes-Gold S, Cohen AE. 2022. Membrane tension propagation couples axon growth and collateral branching. bioRxiv 2022.01.09.475560.

Steinkühler J, Fonda P, Bhatia T, Zhao Z, Leomil FSC, Lipowsky R, Dimova R. 2021. Superelasticity of Plasma- and Synthetic Membranes Resulting from Coupling of Membrane Asymmetry, Curvature, and Lipid Sorting. Adv Sci 8:1–9. doi:10.1002/advs.202102109

Steinkühler J, Sezgin E, Urbančič I, Eggeling C, Dimova R. 2019. Mechanical properties of plasma membrane vesicles correlate with lipid order, viscosity and cell density. Commun Biol 2:1–8. doi:10.1038/s42003-019-0583-3

Sukharev SI, Blount P, Martinac B, Blattnert FR, Kung C. 1994. A large-conductance mechanosensitive channel in E. coli encoded by mscL alone. Nature 368:265–268.

Sukharev SI, Sigurdson WJ, Kung C, Sachs F. 1999. Energetic and spatial parameters for gating of the bacterial large conductance mechanosensitive channel, MscL. J Gen Physiol 113:525–539. doi:10.1085/jgp.113.4.525

Sun W, Chi S, Li Yuheng, Ling S, Tan Y, Xu Y, Jiang F, Li J, Liu C, Zhong G, Cao D, Jin X, Zhao D, Gao X, Liu Z, Xiao B, Li Yingxian. 2019. The mechanosensitive Piezo1 channel is required for bone formation. Elife 8:1–25. doi:10.7554/eLife.47454

Teng J, Loukin S, Anishkin A, Kung C. 2015. The force-from-lipid (FFL) principle of mechanosensitivity, at large and in elements. Pflügers Arch - Eur J Physiol 467:27–37. doi:10.1007/s00424-014-1530-2

Ursell T, Huang KC, Peterson E, Phillips R. 2007. Cooperative gating and spatial organization of membrane proteins through elastic interactions. PLoS Comput Biol 3. doi:10.1371/journal.pcbi.0030081

Vásquez V, Krieg M, Lockhead D, Goodman MB. 2014. Phospholipids that Contain Polyunsaturated Fatty Acids Enhance Neuronal Cell Mechanics and Touch Sensation. Cell Rep 6:70–80. doi:10.1016/j.celrep.2013.12.012

Wang B, Lane BJ, Kapsalis C, Ault JR, Sobott F, Mkami H El, Calabrese AN, Kalli AC, Pliotas C. 2021. Tension mediated mechanical activation and pocket delipidation lead to an analogous MscL state. bioRxiv.

Wang Y, Guo Y, Li G, Liu C, Wang L, Zhang A, Yan Z, Song C. 2021. The push-to-open mechanism of the tethered mechanosensitive ion channel nompc. Elife 10:1–20. doi:10.7554/eLife.58388

Wang Y, Liu Y, Deberg HA, Nomura T, Hoffman MT onks, Rohde PR, Schulten K, Martinac B, Selvin PR. 2014. Single molecule FRET reveals pore size and opening mechanism of a mechano-sensitive ion channel. Elife 3:e01834. doi:10.7554/eLife.01834

Wiggins P, Phillips R. 2005. Membrane-protein interactions in mechanosensitive channels. Biophys J 88:880–902. doi:10.1529/biophysj.104.047431

Wolff U. 2004. Monte Carlo errors with less errors. Comput Phys Commun 156:143–153. doi:10.1016/S0010-4655(03)00467-3

Xian Tao Li, Dyachenko V, Zuzarte M, Putzke C, Preisig-Müller R, Isenberg G, Daut J. 2006. The stretch-activated potassium channel TREK-1 in rat cardiac ventricular muscle. Cardiovasc Res 69:86–97. doi:10.1016/j.cardiores.2005.08.018

Xue F, Cox CD, Bavi N, Rohde PR, Nakayama Y, Martinac B. 2020. Membrane stiffness is one of the key determinants of E. coli MscS channel mechanosensitivity. Biochim Biophys Acta - Biomembr 1862:183203. doi:10.1016/j.bbamem.2020.183203

Yoo J, Cui Q. 2009. Curvature generation and pressure profile modulation in membrane by lysolipids: Insights from coarse-grained simulations. Biophys J 97. doi:10.1016/j.bpj.2009.07.051

Yoshimura K, Batiza A, Kung C. 2001. Chemically Charging the Pore Constriction Opens the Mechanosensitive Channel MscL. Biophys J 80:2198–2206. doi:10.1016/S0006-3495(01)76192-9

Zaki AM, Carbone P. 2017. How the Incorporation of Pluronic Block Copolymers Modulates the Response of Lipid Membranes to Mechanical Stress. Langmuir 33:13284–13294. doi:10.1021/acs.langmuir.7b02244

Zhou Y, Raphael RM. 2005. Effect of salicylate on the elasticity, bending stiffness, and strength of SOPC membranes. Biophys J 89:1789–1801. doi:10.1529/biophysj.104.054510

